# pSNAP: Proteome-wide analysis of elongating nascent polypeptide chains

**DOI:** 10.1101/2021.09.22.461445

**Authors:** Junki Uchiyama, Rohini Roy, Dan Ohtan Wang, Kazuya Morikawa, Yuka Kawahara, Mio Iwasaki, Chiaki Yoshino, Yuichiro Mishima, Yasushi Ishihama, Koshi Imami

## Abstract

Cellular global translation is often measured using ribosome profiling or quantitative mass spectrometry, but these methods do not provide direct information at the level of elongating nascent polypeptide chains (NPCs) and associated co-translational events. Here we describe pSNAP, a method for proteome-wide profiling of NPCs by affinity enrichment of puromycin- and stable isotope-labeled polypeptides. pSNAP does not require ribosome purification and/or chemical labeling, and captures *bona fide* NPCs that characteristically exhibit protein N-terminus-biased positions. We applied pSNAP to evaluate the effect of silmitasertib, a potential molecular therapy for cancer, and revealed acute translational repression through casein kinase II and mTOR pathways. We also characterized modifications on NPCs and demonstrated that the combination of different types of modifications, such as acetylation and phosphorylation in the N-terminal region of histone H1.5, can modulate interactions with ribosome-associated factors. Thus, pSNAP provides a framework for dissecting co-translational regulations on a proteome-wide scale.

## Introduction

Co-translational regulation, such as modifications of nascent polypeptide chains (NPCs) during translation, drives many aspects of cellular proteostasis, including protein folding, processing, subcellular targeting, and translational control (Aviner et al., 2021; Collart and Weiss, 2020; Schwarz and Beck, 2019). Therefore, monitoring co-translational events at the NPC level is crucial for understanding cellular proteome dynamics at the moment a peptide is born.

Current approaches for systematically profiling the newly synthesized proteome are mainly based on the capture of proteins metabolically labeled with stable isotope-labeled (SILAC) amino acids (Doherty et al., 2009; Klann et al., 2020; Schwanhäusser et al., 2009) or bioorthogonal amino acids such as azidohomoalanine (Dieterich et al., 2006; Eichelbaum et al., 2012; McShane et al., 2016). These methods allow us to profile mainly fully translated products (Eichelbaum et al., 2012; McShane et al., 2016), but cannot enrich NPCs actively being elongated by ribosomes in action. In contrast, puromycin is an aminoacylated tRNA analog that can be incorporated at the C-termini of elongating NPCs (Aviner, 2020). Hence, puromycin labeling has been extensively used to monitor protein synthesis in many applications, including imaging and immunoblotting, and in various systems ranging from cell-free translation to cultured cells and whole animals (Aviner, 2020). However, the utility of puromycin or its derivatives for proteome-wide analysis of NPCs has been limited due to the need for complicated procedures, including ribosome purification by ultracentrifugation (Aviner et al., 2013) and/or chemical labeling (Forester et al., 2018; Huang et al., 2021; Tong et al., 2020; Uchiyama et al., 2021) prior to affinity purification of NPCs. Furthermore, reliable detection of NPCs is often hampered by non-specific binding of high-background pre-existing proteins to beads or resin during affinity purification (Eichelbaum et al., 2012; Howden et al., 2013; Mellacheruvu et al., 2013).

To overcome these limitations, we have developed a method that combines quantitative proteomics and dual pulse labeling with puromycin and SILAC amino acids, termed Puromycin- and SILAC labeling-based NAscent Polypeptidome profiling (pSNAP). We demonstrate the broad utility of the method by applying it to both HeLa cells and primary cells to characterize rapid translational changes as well as modifications on NPCs such as protein N-terminal acetylation.

## Results and Discussion

### pSNAP enables global profiling of nascent polypeptide chains

We reasoned that NPCs incorporating puromycin could be immunoprecipitated with an anti-puromycin antibody (**Figure 1A**), thereby allowing for proteome-wide analysis of NPCs by means of liquid chromatography-tandem mass spectrometry (LC/MS/MS). We first confirmed that puromycin incorporation into proteins was translation-dependent (**Figure S1A**) and that no marked degradation of the puromycin-labeled proteins occurred during 2 hr treatment of HeLa cells with 10 μM puromycin (**Figure S1B**). We chose 10 μM puromycin because this concentration could label a wide range of NPCs, while higher concentrations (>30 μM) of puromycin resulted in the production of smaller NPCs (**Figure S1C**). This is consistent with the fact that puromycin binds to NPCs in competition with aminoacyl-tRNAs, and therefore puromycin at very low concentration (0.04 μM) can bind only to full-length NPCs at the C-terminus (Miyamoto-Sato et al., 2000). We then tested a monoclonal antibody against puromycin (clone 12D10) and found that puromycylated proteins could be effectively immunoprecipitated when appropriate amounts of the antibody and input protein were used (**Figure S1D**). Based on these results, we used 15 μg antibodies per 250 μg protein input for subsequent immunoprecipitation (IP) experiments.

**Figure 1.**
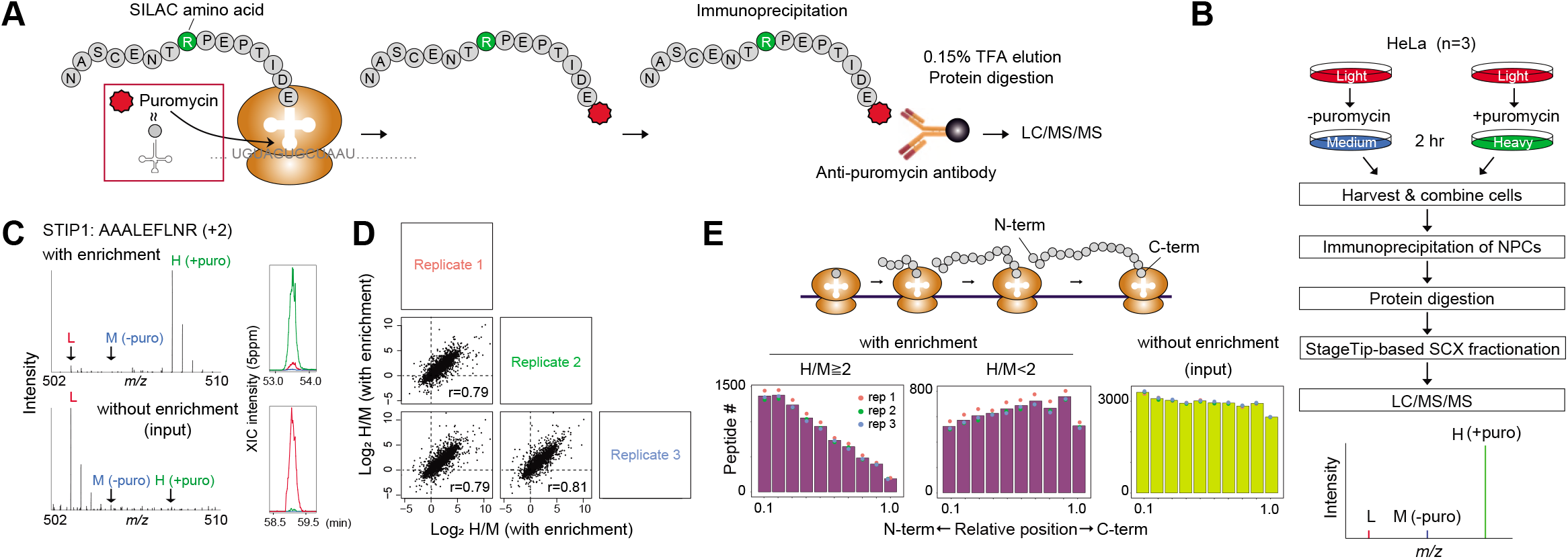
Profiling nascent polypeptide chains by pSNAP. **(A)** Principle of the enrichment of NPCs with the pSNAP. **(B)** Schematic representation of pSNAP workflow. HeLa cells are pulse labeled for 2 hr with a combination of 10 μM puromycin and heavy amino acids or with only medium-heavy amino acids. After IP of NPCs with anti-puromycin antibodies, NPCs are eluted with 0.15% TFA and digested into tryptic peptides. The resulting peptide sample is fractionated into 7 fractions using an SCX StageTip (see Method) and each fraction is analyzed by LC/MS/MS. Heavy-to-medium (H/M) ratios represent the degree of enrichment of NPCs. **(C)** Exemplary MS spectra for AAALEFLNR (STIP1) obtained with (top) and without (bottom) enrichment. The isotope clusters of the L, M, and H peaks correspond to nonspecific proteins from the pre-existing proteome pool and from the ‘medium’-labeled proteins and ‘heavy’-labeled NPCs, respectively. Extracted ion chromatograms (XICs) of monoisotopic peaks of the L, M and H peptides are shown. **(D)** Multi-scatter plots of log_2_ H/M ratios from three independent experiments of pSNAP. **(E)** pSNAP can enrich NPCs. (top) The ribosome elongates an NPC from its N-terminal end to the C-terminal end. Thus, positions of NPC-derived tryptic peptides are biased towards the N-termini of proteins. (bottom) Relative starting positions of identified peptides within proteins. The bars represent averaged values from three independent experiments.

We next sought to profile individual NPCs with LC/MS/MS. For proof-of-concept, HeLa cells were pulse-labeled with a combination of puromycin and ‘heavy (H)’ amino acids (Arg’10’ and Lys’8’) for 2 hr (**Figure 1B**). As a control, cells were treated with only ‘medium-heavy (M)’ amino acids (Arg’4’ and Lys’4’) and puromycin was omitted. The use of ‘M’ and ‘H’ labeling enabled us to distinguish *bona fide* NPCs (H-labeled) from non-specific proteins [light (L)- or M-labeled]. As expected, H-labeled proteins were highly enriched in the IP sample (**Figure 1C top**) while preexisting proteins were predominantly observed in the input (**Figure 1C bottom**). Overall, we observed high H/M ratios (≧2) for 70% of 2,619 quantified proteins with good reproducibility (**Figure 1D, Figure S1E** and **Table S1**), demonstrating that pSNAP can profile thousands of NPCs. In contrast, we observed low H/L ratios (<1) for 62 % of the quantified proteins (**Figure S 1F**) due to the high background resulting from non-specific binding of pre-existing proteins (L) to beads. These results highlight the importance of pulse labeling with SILAC amino acids to differentiate NPCs and pre-existing proteins.

### Proteins captured by pSNAP exhibit a signature of elongating nascent polypeptides

Ribosomes elongate NPCs from their amino (N-) terminal end to their carboxy (C-) terminal end (**Figure 1E top**). In line with this directionality, the mapped positions of peptides with H/M≧2 clearly showed a bias towards the N-termini of the corresponding proteins (**Figure 1E bottom**), further supporting the enrichment of elongating NPCs. Notably, such a trend was not observed for less enriched peptides (i.e., H/M<2) or input cell lysates (**Figure 1E bottom**).

To further evaluate pSNAP, we compared it with two orthogonal approaches, pSILAC (Schwanhäusser et al., 2009) and ribo-seq (Ingolia et al., 2009), used for studying protein synthesis at the levels of protein and mRNA, respectively. To this end, we used our input data (**Figure 1E**) as a 2 hr pSILAC experiment and a publicly available ribo-seq dataset for HeLa cells (Stumpf et al., 2013). We found moderate correlations with the two independent methods; r = 0.49 (vs. pSILAC) and r = 0.40 (vs. ribo-seq) (**Figure S1G**); similar levels of correlation were also seen in a previous study using a biotin-puromycin-based approach (Aviner et al. Gene. Dev. 2014). These results indicate that pSNAP can accurately quantify translation products.

### pSNAP with short pulse labeling enables quantitative profiling of nascent proteome

We next assessed the effect of pulse-labeling time on NPC capture. Previous studies using O-propargyl-puromycin (OPP) (Forester et al., 2018; Uchiyama et al., 2021) employed 2 hr labeling, while a more recent study (Tong et al., 2020) performed only 15 min labeling. However, the trend of protein N-terminus-biased positions was not seen in the 15 min labeling method, in contrast to our results (**Figure 1E bottom left**), implying that NPCs cannot be sufficiently captured with such a short labeling time, or are hidden by non-specific binding of pre-existing proteins. Based on these prior experiments and the immunoblotting results (**Figure S1B left panel**), we chose 30 min as the shortest effective labeling time and used increased protein inputs (250, 500, and 750 μg) for pSNAP experiments. We found that this labeling time enabled us to profile NPCs. We observed high H/M ratios (**Figure S2A, Table S2**) and protein N-terminus-biased positions (**Figure S2B**), as seen in the case of 2 hr labeling (**Figure 1E left panel**), though slightly fewer NPCs were quantified in the short labeling method compared to 2 hr labeling (**Table S2**).

To further demonstrate the validity of the 30 min labeling method, we analyzed lipopolysaccharide (LPS)-induced translational responses with different periods of pulse labeling (30 min, 1 hr, and 2 hr). RAW264.7 macrophages were first treated with 100 ng/mL LPS for 1 hr, and then switched to a medium containing puromycin and heavy amino acids. As a control, cells were treated with vehicle (water), and pulse-labeled with puromycin and medium-heavy amino acids. We successfully quantified NPCs including known LPS-regulated genes (e.g., Cxcl2, Junb, Rela, and Nfkbiz) regardless of the pulse labeling time (**Figures S2C and S2E, Table S2**). These results established that even a labeling time as short as 30 min can be used to robustly detect translational responses using the pSNAP approach. It should be noted that we used a 2 hr labeling time for the following experiments because it did not significantly affect cell viability (**Figure S2D**) or quantitative analysis (**Figure S2C**) and yielded sufficient amount of puromycylated NPCs (**Table S2**).

### pSNAP allows quantitative nascent proteome profiling in primary neuronal cultures

In addition to HeLa cells, we applied pSNAP to mouse primary cortical neuron cultures, which contain a mixture of brain cell types but are highly enriched for neurons (**Figure 2E**). pSNAP revealed strong enrichment of peptides near the N-termini of proteins (**Figures 2A-C**), and achieved highly sensitive detection of NPCs, including neuronal markers; NCAM1, FOXG1, and DCX (**Table S3**), supporting the validity of NPCs detection using pSNAP. We also quantified the differential NPC profiles between 5 and 14 days in vitro (DIV), and identified 122 and 203 significantly up-regulated proteins in DIV5 and DIV14, respectively. (**Figures 2D-F** and **Table S3**). These results overall reflect the known proteome dynamics during neuronal development in vitro (Frese et al., 2017). For example, nascent proteins related to ‘synaptic vesicle cycle’ were overrepresented in DIV14 (**Figure 2G**), consistent with the occurrence of synapse formation and maintenance after development. In addition, both immunostaining and pSNAP confirmed that a presynaptic marker synaptophysin (SYPH) was upregulated in DIV14 (**Figure 2E**). Collectively, these results indicate that pSNAP captures genuine dynamics of NPCs in primary cultures and enables quantitative nascent proteome analysis.

**Figure 2.**
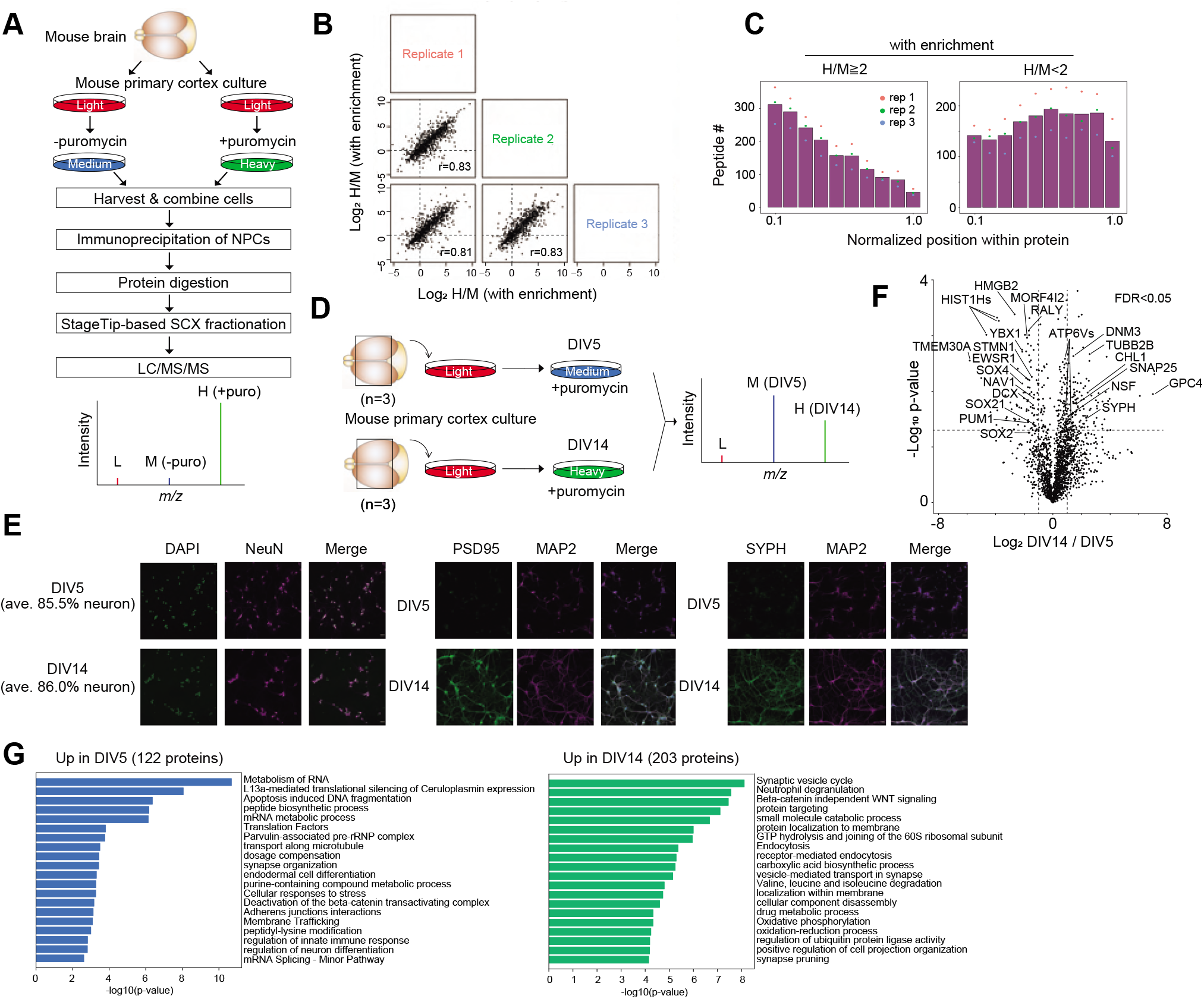
Application of pSNAP to mouse primary cortical neuronal cultures. (A) Schematic representation of the pSNAP workflow. Mouse primary cortical neurons are pulse labeled for 2 hr with a combination of 10 μM puromycin and heavy amino acids or only mediumheavy amino acids. After IP of NPCs with anti-puromycin antibodies, NPCs are eluted with 0.15% TFA and digested into tryptic peptides. The resulting peptide sample is analyzed by LC/MS/MS. (B) Multi-scatter plots of log_2_ H/M ratios from three independent experiments using mouse primary neurons. **(C)** pSNAP can enrich NPCs from primary neurons. Relative starting positions of identified peptides within proteins. The bars represent averaged values from three independent experiments. **(D)** Experimental design for the differential nascent proteome profiling of the mouse primary cultures between DIV 5 and 14. **(E)** Representative images of primary neuronal cultures (DIV5 and DIV14) stained with DAPI (nuclear marker), NeuN (neuronal marker), MAP2 (dendritic microtubule marker), PSD95 (postsynaptic marker) and SYPH (presynaptic marker). Scale bar, 30 μm. **(F)** A volcano plot showing differential NPC levels between DIV 5 and 14. Significantly regulated proteins were identified based on a combination of the t-test p-value (p<0.05) of three replicates and the mean of the SILAC ratios [above 0.5 (a log_2_ ratio)] (dash lines) which correspond to FDR <0.05 (see Method). **(G)** Gene ontology enrichment analyses for the significantly up-regulated NPCs (adjusted p < 0.01).

### Quantifying translational responses induced by a kinase inhibition

The ability to capture and quantify NPCs proteome-wide with high accuracy enables quantitative measurements of acute translational changes that allow cells to respond to specific stimuli. To illustrate this, we next applied pSNAP to characterize the global impact of a kinase inhibitor on cellular translation. We focused on casein kinase 2 (CK2) as it is ubiquitously expressed in all cells, and has been implicated in translational control through phosphorylation of specific eukaryotic initiation factors (eIFs) (Gandin et al., 2016; Lamper et al., 2020). However, the protein targets of translational regulation by CK2 remain unknown. Therefore, HeLa cells were preincubated with either DMSO or a specific CK2 inhibitor silmitasertib (also known as CX4945) (Siddiqui-Jain et al., 2010) for 10 min (**Figure 3A**), and then pulse-labeled with puromycin and SILAC amino acids for 2 hr in the presence of DMSO or silmitasertib, and processed as illustrated in **Figure 1B**. H/M ratios in MS spectra represent the difference in production of NPCs between the two conditions (silmitasertib and DMSO treatments). We first confirmed that NPCs could be enriched; H- and M-labeled peptides both exhibited the trend of protein N-terminus-biased positions (**Figure S3A**).

**Figure 3.**
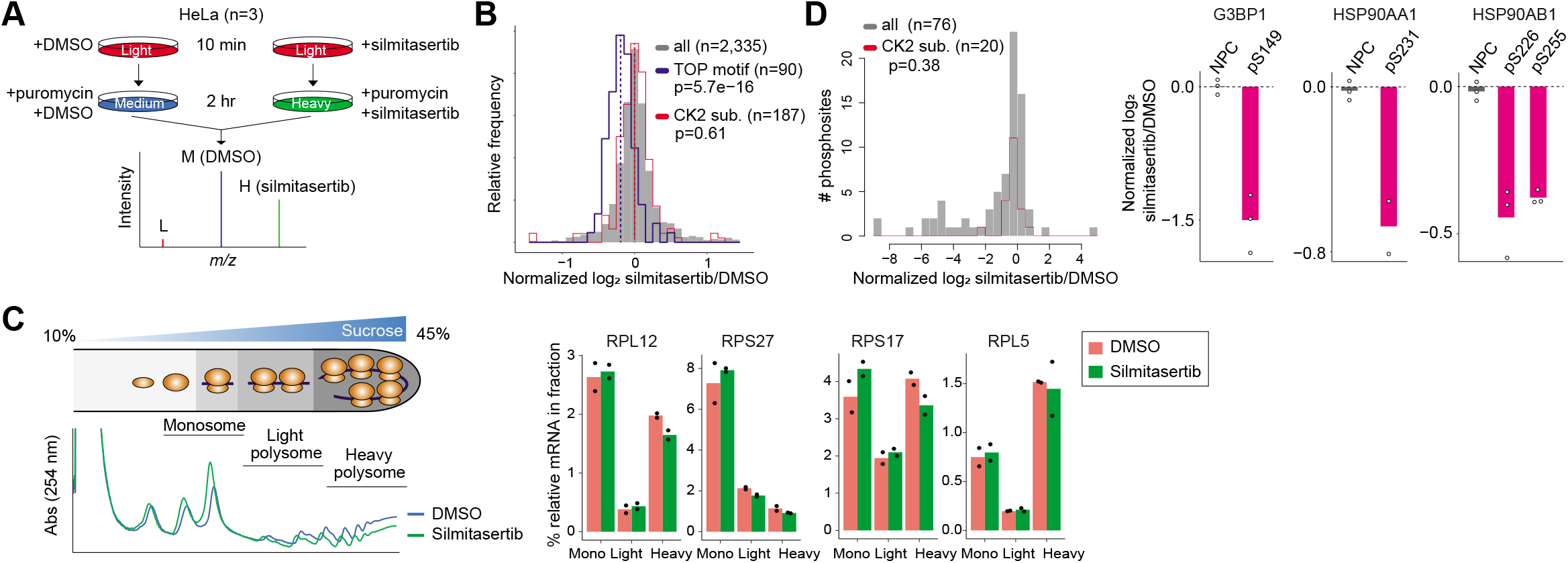
Quantitative analysis of translational responses and nascent phosphorylation. **(A)** Experimental design for global analysis of translational changes induced by a CK2 inhibitor silmitasertib (10 μM). **(B)** A histogram of log_2_ fold changes [H/M (silmitasertib/DMSO) ratios] of protein synthesis induced by silmitasertib. Averaged log_2_ H/M ratios based on proteins quantified in at least two of the three replicates are shown. Results from individual replicates are shown in **Figure 3B**. A subset of proteins whose mRNAs contain a TOP motif is shown in dark blue. CK2 substrates determined by an in vitro kinase reaction (Sugiyama et al., 2019) are shown in red. All quantified proteins are shown in grey. The dashed lines indicate median values of the three groups (all, TOP motif, and CK2 substrates) The p-value was computed using the two-sided Wilcoxon rank-sum test. **(C)** qRT-PCR validation of pSNAP result. (Left panel) Representative polysome profiles of HeLa cells treated with DMSO or silmitasertib. (Right panel) Polysomal association of mRNAs encoding for the selected ribosomal proteins that were repressed upon CK2 inhibition according to pSNAP. **(D)** CK2 inhibition led to decreased phosphorylation levels of nascent forms of known CK2 substrates. (Left panel) A histogram of log_2_ fold changes [H/M (silmitasertib/DMSO) ratios] of phosphorylation sites induced by silmitasertib. Averaged log_2_ H/M ratios based on class I phosphosites quantified in at least one of the three replicates are shown. CK2 substrates determined by in vitro kinase reaction (Sugiyama et al., 2019) are shown in red. All quantified proteins are shown in grey. The p-value was computed using the two-sided Wilcoxon rank-sum test. (Right panel) Change of NPC and phosphorylation levels due to silmitasertib treatment versus DMSO treatment. The indicated phosphorylation sites of G3BP1, HSP90AA1 and HSP90AB1 are known CK2 substrates. Levels of the NPCs were quantified from NPC-derived unphosphorylated peptides. The bars represent averaged values from two or three replicates.

To understand the gene expression networks at the NPC level, we first asked whether in vitro CK2 substrates (Sugiyama et al., 2019) might be regulated translationally, but found no evidence to support this (**Figure 3B, Table S4**). On the one hand, a recent study showed that CK2 acts in concert with mTORC1 (Gandin et al., 2016) and regulates the translation of mRNAs containing 5’ terminal oligopyrimidine (TOP) motifs (Hsieh et al., 2012; Thoreen et al., 2012). We therefore asked whether the inhibition of CK2 affects the translation of TOP mRNAs. Indeed, we found that CK2 inhibition led to marked repression of NPCs with a TOP motif in their mRNAs (p = 5.7e-16), encoding for components of the translational machinery such as ribosomal proteins (**Figure 3B** and **Figure S3B** for individual replicates).

To validate this, we performed quantitative real-time qRT-PCR analysis of mRNAs encoding for the selected silmitasertib-sensitive ribosomal proteins (average log_2_ fold-change, RPL12: −0.52, RPS27: −0.54, RPS17: −0.16, RPL5: −0.18) from monosome, light and heavy polysome fractions (**Figure 3C left panel**). The polysome profiles revealed that silmitasertib induced a slight reduction in the polysomes (**Figure 3C left panel**), indicating that CK2 inhibition leads to translational repression. The silmitasertib-sensitive group (RPL12, RPS27, and RPS17 but not RPL5) tended to be less abundant in the heavy polysome fraction than in the monosome fraction (**Figure 3C right panel**), in agreement with the pSNAP result (**Figure 3B**). Although qRT-PCR confirmed the general trend in translational regulation revealed by pSNAP, there was no correlation between the effect sizes measured with pSNAP and those observed with qRT-PCR. This may be due to technical differences between methods (mass spectrometry or PCR) and analytes [elongating nascent polypeptides (pSNAP) or ribosome-associated mRNA (qRT-PCR)]. TOP mRNAs are well-known targets that are subject to selective translation through mTORC1 (Hsieh et al., 2012; Thoreen et al., 2012). Hence, this result supports the idea that CK2 may regulate the translation of TOP mRNAs in concert with mTORC1, in line with a previous report that CK2 enhances mTORC1 activity (Gandin et al., 2016). In sum, our analysis uncovered acute translational responses to silmitasertib via the CK2 and mTOR pathways, which may contribute to further understanding of the mechanism of action of silmitasertib, a promising molecular therapy for several types of cancers (phase II) (Siddiqui-Jain et al., 2010) as well as SARS-CoV-2 infection (Bouhaddou et al., 2020).

### Profiling modifications on nascent proteins

The present method not only enables the global profiling of NPCs, but also highlights the modifications of NPCs that might be co-translationally regulated. Conventional proteomic approaches cannot resolve modifications on nascent and matured proteins, and so the distribution of the two types of modifications within a protein goes undetected; yet the timing of modifications can be important for protein processing (Aksnes et al., 2019; Varland et al., 2015) such as folding (Keshwani et al., 2012; Kii et al., 2016). In the silmitasertib treatment experiment (**Figure 3**), we identified 127 unique phosphopeptides without phosphopeptide enrichment. Among them, 76 class I phosphosites (localization probability >0.75) (Olsen et al., 2006) that were quantified in at least one of the three replicates are shown in **Figure 3D**. We found that CK2 inhibition led to decreased phosphorylation levels on nascent forms of known CK2 substrates, such as G3BP1 pS149 (Reineke et al., 2017), HSP90AA1 pS231, and HSP90AB1 pS226, pS255 (Mollapour and Neckers, 2012), while no marked change was observed at the NPC level (**Figure 3D right panel** and **Table S4**). On the one hand, we found that phosphorylation of most of the CK2 substrates was not reduced by the inhibitor (**Figure 3D left panel**), indicating that phosphorylation of NPCs may be differentially regulated from that of mature proteins. Hence, CK2 may act in close proximity to the ribosome to co-translationally phosphorylate specific nascent proteins, possibly regulating protein stability through phosphorylation of newly made proteins, as observed for XRCC1 (Parsons et al., 2010) and CFTR (Pankow et al., 2019).

Protein N-terminal (Nt) acetylation is one well-studied ‘co-translational’ modification (Aksnes et al., 2019; Yeom et al., 2017); however, recent studies have revealed ‘post-translational’ Nt-acetylation on many transmembrane proteins and actin (Yeom et al., 2017). While earlier N-terminomics studies identified thousands of protein Nt-acetylation sites (Choudhary et al., 2014; Lai et al., 2015; Yeom et al., 2017), it remains unclear whether these sites are co- or post-translationally modified. We thus sought to apply pSNAP to pinpoint Nt-acetylation sites on NPCs with high accuracy. For this purpose, HeLa cells were treated with either 100 μg/mL cycloheximide (CHX) or DMSO for 2 hr in the presence of 10 μM puromycin and corresponding SILAC amino acid pairs. By combining pSNAP with low pH strong cation exchange (SCX) chromatography (Helbig et al., 2010), we enriched Nt-acetylated peptides that were eluted in the flow-through fraction and early in SCX fractionation due to the loss of positive charge at their N-terminal ends (**Figure 4A**). We confirmed that the NPCs could be enriched (**Figure S4** and **Table S5**) and identified 298 unique protein Nt-acetylated sites that exhibited H/M (DMSO/CHX) ≧2 in at least one of the three replicates (**Table S5**). Notably, beta-actin’s Nt-acetylation showed H/M<1 in all replicates, indicating that it occurs post-translationally, in agreement with a previous report (Drazic et al., 2018). To better understand acceptor sites for Nt-acetylation on NPCs, we focused on amino acids at the second residue [next to the initiator methionine (iMet)] (**Figure 4B**). In accordance with the substrate specificity of major N-terminal acetyltransferases (Aksnes et al., 2019; Yeom et al., 2017) and known Nt-acetylation sites from the Uniprot human database, we observed a high prevalence of alanine, serine, and threonine for Nt-acetylated NPCs whose iMet was cleaved, while the acidic amino acids (aspartic acid and glutamic acid) and phenylalanine were overrepresented in the iMet-retained and Nt-acetylated NPCs (**Figure 4B**). We did not observe a significant difference in acceptor amino acids between NPCs and the input.

**Figure 4.**
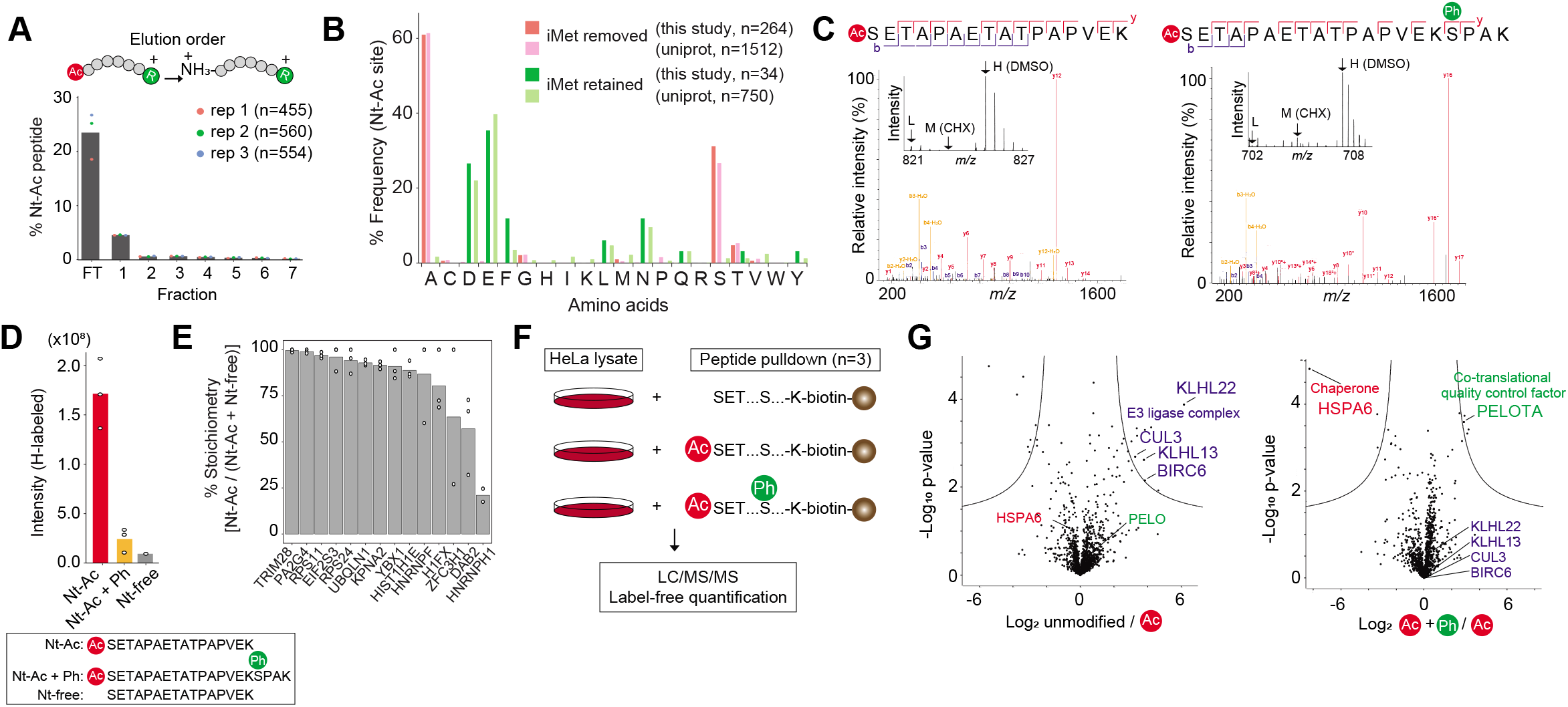
Characterization of protein Nt-acetylation and phosphorylation on nascent proteins. **(A)** Enrichment of Nt-acetylated peptides with SCX-based fractionation. The percentage of the number of protein Nt-acetylated peptides in all peptides identified in individual fractions is shown. The bars represent averaged values from three replicates. **(B)** The amino acid frequency at the second residue next to iMet of acetylated protein N-termini based on the absence (pink) or presence (green) of iMet. Only nascent proteins that showed H/M>2 in at least one of the three replicates were considered. For comparison, the human proteome from the SwissProt protein database is shown (light pink and light green indicate iMet-removed and iMet-retained sites, respectively). **(C)** Representative MS/MS spectra of Ac-SETAPAETATPAPVEK (left) and Ac-SETAPAETATPAPVEKpSPAK (right) from histone H1.5. The insets show representative MS spectra of corresponding peptides and demonstrate that the protein Nt-acetylation and adjacent phosphorylation occurred on nascent H1.5. **(D)** Quantification of H-labeled peptide intensities from Nt-acetylated (red), Nt-acetylated and phosphorylated (orange), and Nt-unmodified (grey) forms. **(E)** Stoichiometry (%) of protein Nt-acetylation estimated based on intensities of H-labeled peptides from Nt-acetylated and counterpart (unmodified) forms. The bars represent averaged values from two or three replicates. **(F)** Experimental design for peptide-pulldown assays using three different peptide probes. Nt-unmodified, Nt-acetylated, or Nt-acetylated and phosphorylated biotinylated peptides corresponding to amino acid residues from 2 to 22 (SETAPAETATPAPVEKSPAKK) of H1.5 were conjugated to streptavidin agarose resins. Beads were incubated with HeLa cell lysate, and eluted for LC/MS/MS analysis. **(G)** Volcano plots from the pulldown assays of Nt-free vs Nt-acetylated peptides (left) and Nt-acetylated vs Nt-acetylated and phosphorylated peptides (right) are shown. The cutoff curve indicates false discovery rate (FDR)<0.05, S0>2.

### Protein N-terminal modifications on a nascent protein can modulate binding partners

Protein N-termini are hotspots for modifications during translation and thus can regulate co-translational events such as folding and degradation through interactions with proteins (Collart and Weiss, 2020). We next sought to discover cross-talk between protein Nt-acetylation and other modifications on NPCs as the combination of different types of modifications confers additional specificity and combinatorial logic to protein interactions. We searched our dataset focusing on phosphorylation, which can function as a versatile switch to modulate protein interactions. We identified 134 phosphorylated peptides in the Nt-acetylation-enriched samples. Among them, we focused on phosphorylation at Ser19 of histone H1.5 (H1.5) co-occurring with protein Nt-acetylation within the same peptide (**Figure 4C**). Peptide-level quantification of H-labeled peptides indicated that the Nt-acetylated form (Ac-SETAPAETATPAPVEK) is a major nascent proteoform of H1.5 in comparison to the Nt-acetylated and phosphorylated form (Ac-SETAPAETATPAPVEKpSPAK) and the unmodified form (SETAPAETATPAPVEK) (**Figure 4D**). Because the trypsin cleavage site (Lys18) is next to the phosphorylation site (Ser19), the phosphorylated form was exclusively observed as a missed cleavage peptide due to the inaccessibility of trypsin. We therefore used the no-missed-cleavage peptide (SETAPAETATPAPVEK) as an unmodified counterpart for the “missed-cleavage phosphopeptide” (Ac-SETAPAETATPAPVEKpSPAK) in the quantification shown in **Figure 4D**. Such highly stoichiometric patterns of nascent Nt-acetylation were also seen for the 13 sites whose Nt-acetylated and counterpart (unmodified) peptides were both quantified (**Figure 4E**), in marked contrast to the very low (median 0.02%) stoichiometry of lysine acetylation in HeLa cells (Hansen et al., 2019).

To understand the role of the nascent modifications and their impacts on protein interactions, we performed peptide-based pulldown experiments on HeLa cell lysate using three distinct peptide probes that mimic 1) Nt-unmodified, 2) Nt-acetylated, or 3) Nt-acetylated and phosphorylated forms of H1.5 (**Figure 4F**). The peptide-based screen revealed proteins that differentially interacted with the specific peptide probes (**Figure 4G** and **Table S6**). One prominent example is a ubiquitin E3 ligase complex (KLHL13, KLHL22, CUL3, BIRC6) that showed a robust interaction with the unmodified peptides (**Figure 4G left panel**). Thus, Nt-unmodified H1.5 is likely to be degraded through the ubiquitin-proteasome system, which may explain why the Nt-unmodified H1.5 was markedly less abundant than the acetylated forms in HeLa cells (**Figure 4D**). Accordingly, protein Nt-acetylation of H1.3 is protective against protein degradation, in line with the idea that Nt-acetylated mitochondrial proteins bearing inhibitor of apoptosis binding (IAP) motifs are shielded from the IAP family of E3 ubiquitin ligases (Mueller et al., 2021). Interestingly, the Nt-acetylated and phosphorylated version of the peptide preferentially bound a co-translational quality control factor PELOTA while disfavoring the interaction with a molecular chaperone HSPA6 (**Figure 4G right panel**). PELOTA was shown to promote the dissociation of stalled ribosomes and the release of intact peptidyl-tRNA for ribosome recycling (Pisareva et al., 2011); Thus, the nascent H1.5 phosphorylation in the N-terminal region may represent an additional ‘modification code’ to recruit PELOTA and to repel HSPA6 as a surveillance mechanism for aberrant nascent H1.5. In summary, pSNAP enables us to uncover modifications on NPCs that may represent a new layer of translational control, i.e., one shaped by nascent protein modifications.

## Limitations of the study

While puromycin or its analogue has been used to analyze protein synthesis in many cell lines and model systems (Aviner, 2020), it may also cause a secondary effect on cellular translation (Marciano et al., 2018). In this study, we therefore used a relatively low concentration of puromycin that did not affect cell viability or degradation of puromycylated proteins, at least in HeLa cells, but further investigations would be required to characterize the mode of action of puromycin in detail. We demonstrated that pSNAP is readily applicable to a cell line and to primary-cultured cells, though further development will be needed for its application to in vivo systems. For example, the direct enrichment and proteomic analysis of puromycylated peptides digested from NPCs provides a signature of genuine NPCs, and would not require pulse SILAC labeling.

While we successfully characterized protein Nt-acetylation and phosphorylation at the NPC level, we cannot rule out the possibility of modifications to the released puromycylated NPCs from ribosomes. Further structural analysis and co-localization studies may provide further clarification. Also, since the amount of NPCs obtained by pSNAP is limited (considered to be < 1 μg based on the total ion chromatogram), the method cannot readily be combined with a biochemical enrichment technique for modified peptides. Further improvement of the current pSNAP protocol to increase sensitivity is desirable for capturing omics-level modification sites. As shown in **Figure 3C**, perturbation of cells may cause global changes in translation. However, such global changes would not be detected in a typical proteome analysis because ratios are normalized based on the assumption that abundance of most proteins remains unchanged under two conditions. We also detected M-channel-derived signals for some proteins even without puromycin (**Figures 1D and 2B**). Hence, background noise due to non-specific proteins may compress quantitative ratios in experiments comparing two conditions based on H/M ratios (see e.g., **Figures 2D and 3A**), leading to systematic underestimation of quantitative ratios.

## Conclusions

The pSNAP approach presented here offers multiple advantages over currently available methods for capturing nascent polypeptide chains and their modifications, which represent a hidden layer in understanding translational regulations in cell biology that is inaccessible by conventional ribosome profiling or proteomic approaches. The advantage of pSNAP lies in the use of dual pulse labeling. The incorporation of puromycin facilitates the enrichment of NPCs from a complex background, and the use of pulsed SILAC enables both protein quantification and discrimination of nascent from pre-existing proteins. Moreover, the experimental workflow is simple in contrast to existing methods that involve ribosome purification using ultracentrifugation(Aviner et al., 2013) and/or chemical labeling steps (Forester et al., 2018; Huang et al., 2021; Tong et al., 2020; Uchiyama et al., 2021). In addition, the method does not require special puromycin derivatives such as biotin-puromycin (Aviner et al., 2013) or clickable puromycin (Forester et al., 2018; Huang et al., 2021; Tong et al., 2020; Uchiyama et al., 2021). Our results show that pSNAP can quantify changes in NPC levels in response to environmental cues, and is useful for characterizing nascent modifications. In addition, this method could be applied to identify the NPC interactome during translation since some proteins appear to form homo- (Bertolini et al., 2021) or hetero- (Kamenova et al., 2019) complexes in a co-translational manner.

---

**Figure S1.**
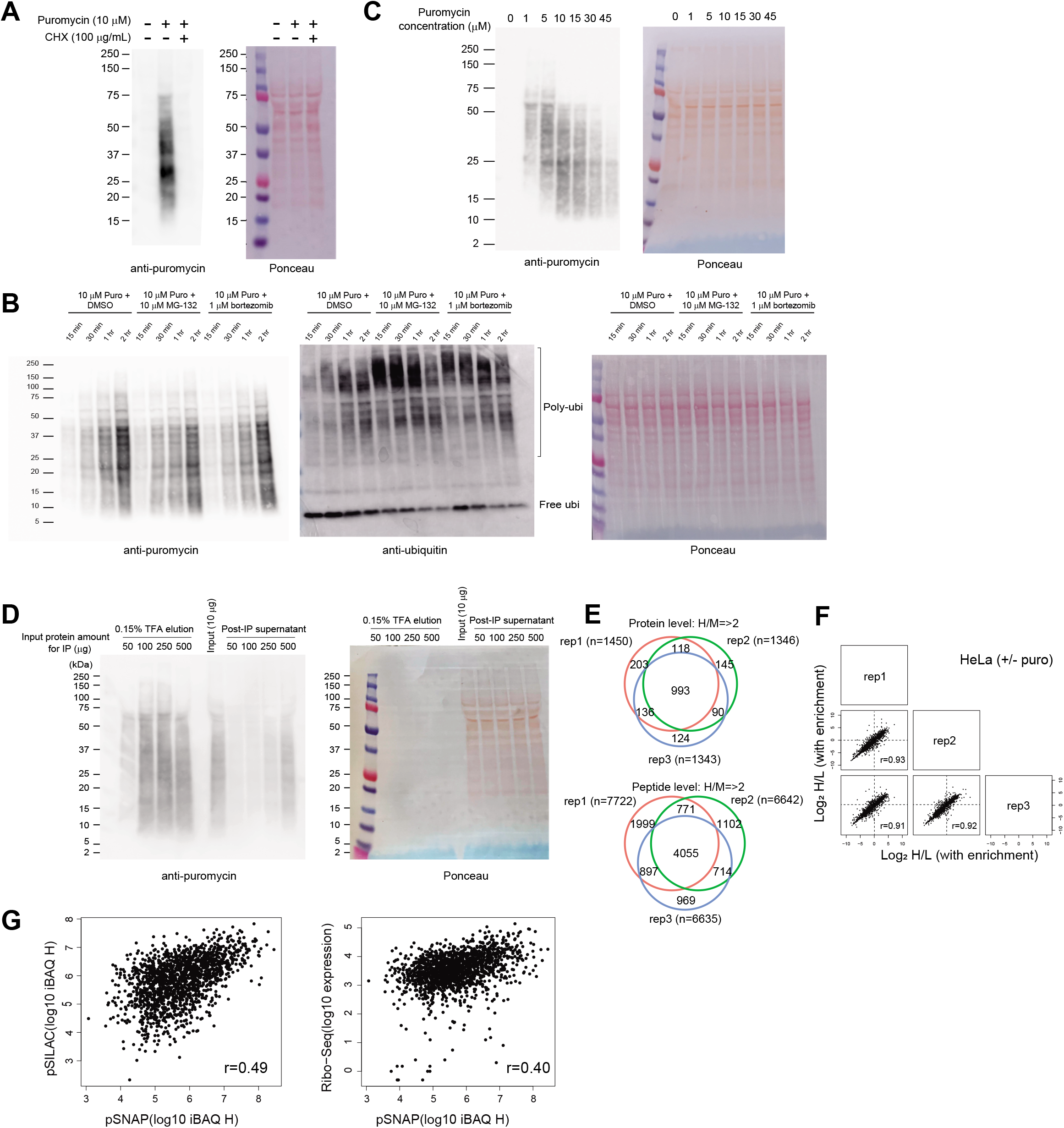
Assessment of puromycin labeling and immunoprecipitation of puromycylated proteins. **(A)** Puromycin labeling of proteins is translation-dependent. Western blot (left) and Ponceau staining (right) of whole-cell lysates of HeLa cells treated with 10 μM puromycin for 2 hr, or 10 μM puromycin and CHX for 2 hr, or neither. **(B)** Abundance of puromycylated proteins was not altered in the presence of a proteasome inhibitor (10 μM MG132 or 1 μM bortezomib) over 2 hr treatment. Western blotting was performed with anti-puromycin antibody (left), anti-ubiquitin antibody (middle), and Ponceau staining (right). This result indicates that puromycylated NPCs were not significantly degraded via the ubiquitin-proteasome system in this time frame, while levels of ubiquitinated total cellular proteins were increased in the presence of proteasome inhibitors. **(C)** Effect of puromycin concentration on production of NPCs. HeLa cells were treated with different concentrations (0, 1, 5, 10, 15, 30, and 45 μM) of puromycin for 2 hr. **(D)** Puromycylated proteins can be immunoprecipitated. Immunoprecipitation (IP) was performed for different protein inputs (50, 100, 250, and 500 μg) with a fixed amount of 15 μg anti-puromycin antibody. The proteins eluted with 0.15% TFA and the remaining supernatant in a post-IP sample were analyzed by means of immunoblotting with the anti-puromycin antibody. 50-250 μg protein inputs were found to be suitable for immunoprecipitation with 15 μg anti-puromycin antibodies. **(E)** Venn diagrams of identified proteins (top) and peptides (bottom) with pSNAP from three independent experiments (see Figure 1). **(F)** Multi-scatter plots of log_2_ H/L ratios from three independent experiments of puromycin (H) and no-puromycin (M) treatments. **(G)** Scatter plots showing the correlation between pSNAP and pSILAC (left) and between pSNAP and ribo-seq (right). The ribo-seq data was taken from a previous study (Stumpf et al., 2013).

**Figure S2.**
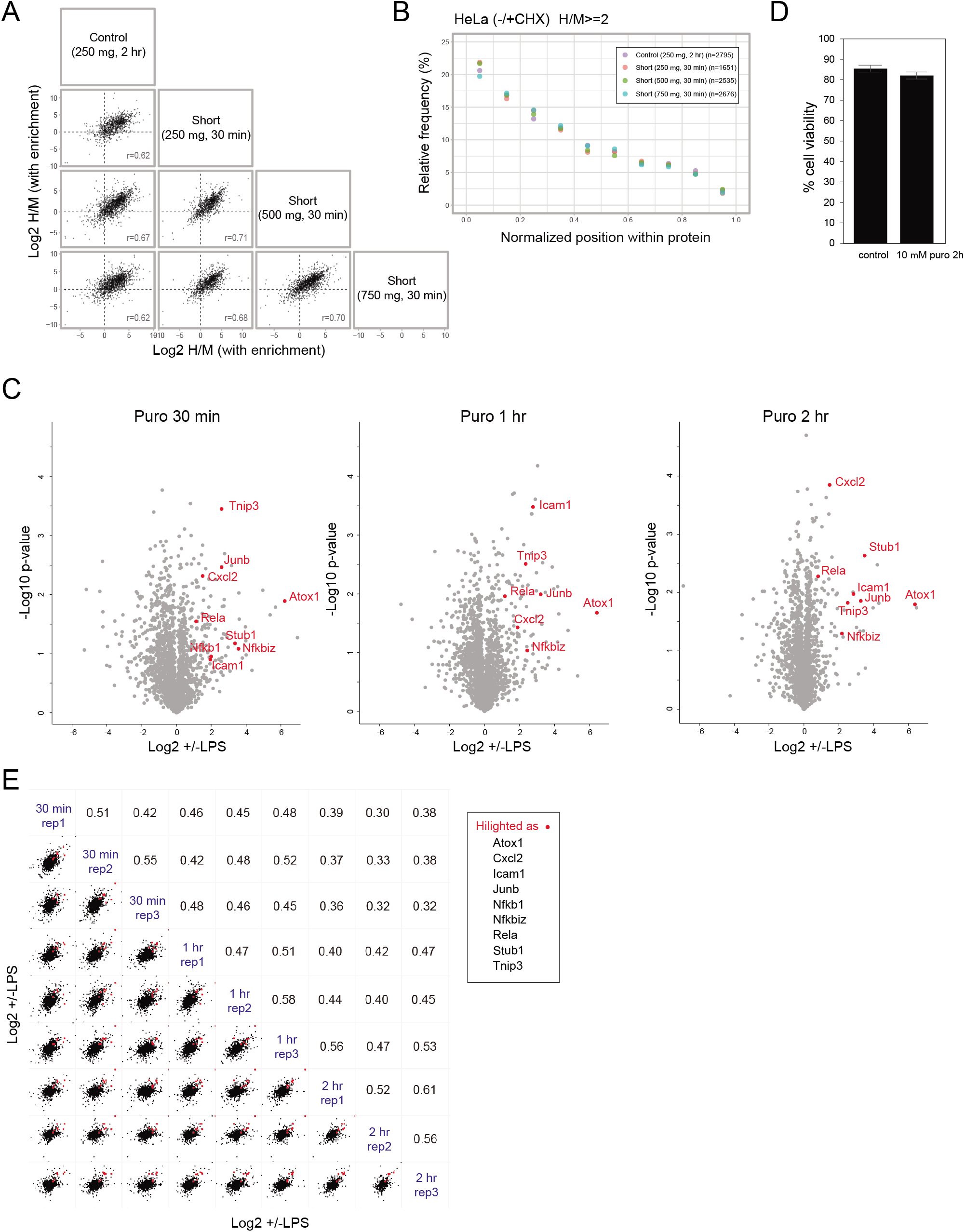
Comparison of puromycin labeling time (30 min vs. 2 hr) in pSNAP. **(A)** Multi-scatter plots of log_2_ H/M ratios from three independent experiments involving 10 μM puromycin (H) and 10 μM puromycin plus 100 μg/mL CHX (M) treatments. 250, 500, or 750 μg protein from HeLa cell lysate was used for the experiment with 30 min pulse labeling, while 250 μg protein was used for 2 hr pulse labeling as a control. **(B)** Relative starting positions of identified peptides within proteins. **(C)** Volcano plots showing LPS-induced translational responses with different lengths of pulse-labeling time (30 min, 1 hr, and 2 hr). RAW264.7 macrophages were first treated with vehicle (water) or 100 ng/mL LPS for 1 hr, and then pulse labeled with 10 μM puromycin and either medium-heavy or heavy amino acids for 30 min, 1 hr or 2 hr. Note that different protein inputs (560, 280, and 140 μg) were used for different pulse labeling times (30 min, 1_hr, and 2 hr, respectively) to obtain sufficient amounts of puromycylated NPCs. **(D)** Trypan blue-based cell viability assay for HeLa cells treated with vehicle (PBS) control or 10 μM puromycin for 2 hr. The bars represent averaged values from three independent experiments. **(E)** Multi-scatter plot showing correlations between log_2_ H/M ratios from three independent experiments at each pulse labeling time (30 min, 1 hr, and 2 hr).

**Figure S3.**
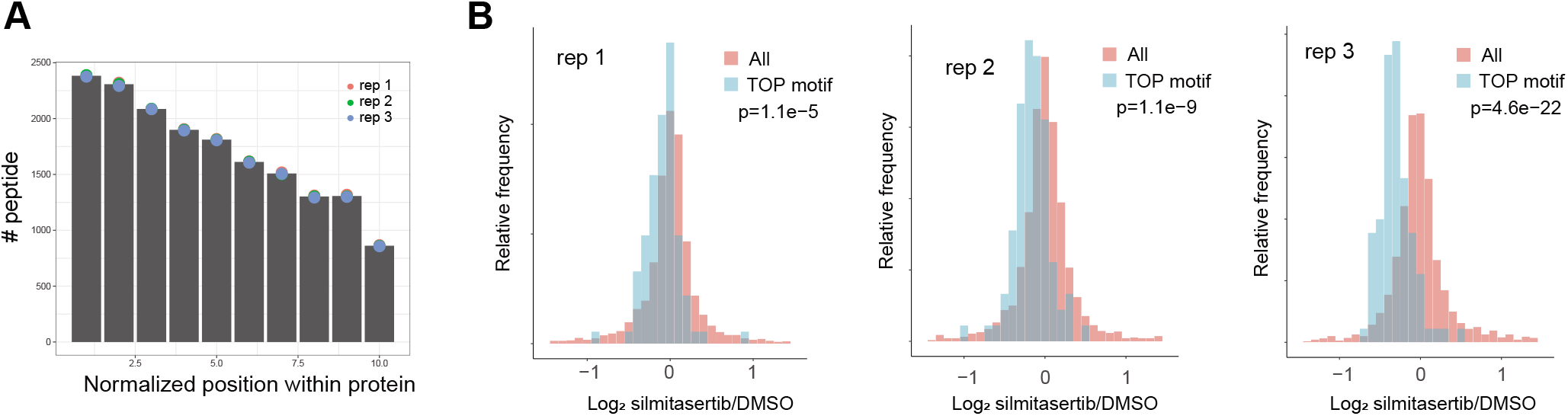
Quantitative analysis of translational changes associated with silmitasertib treatment. **(A)** pSNAP can enrich NPCs from HeLa cells treated with DMSO or silmitasertib. Relative starting positions of identified peptides (only M- or H-labeled) within proteins for NPC-enriched samples. The bars represent averaged values from three independent experiments. **(B)** Histogram of log_2_ FCs [H/M (silmitasertib/DMSO) ratios] of protein synthesis induced by silmitasertib. Results from individual replicates are shown. A subset of proteins whose mRNAs contain a TOP motif is shown in light blue. All quantified proteins are shown in pink. The p values were computed using twosided Wilcoxon rank-sum test.

**Figure S4.**
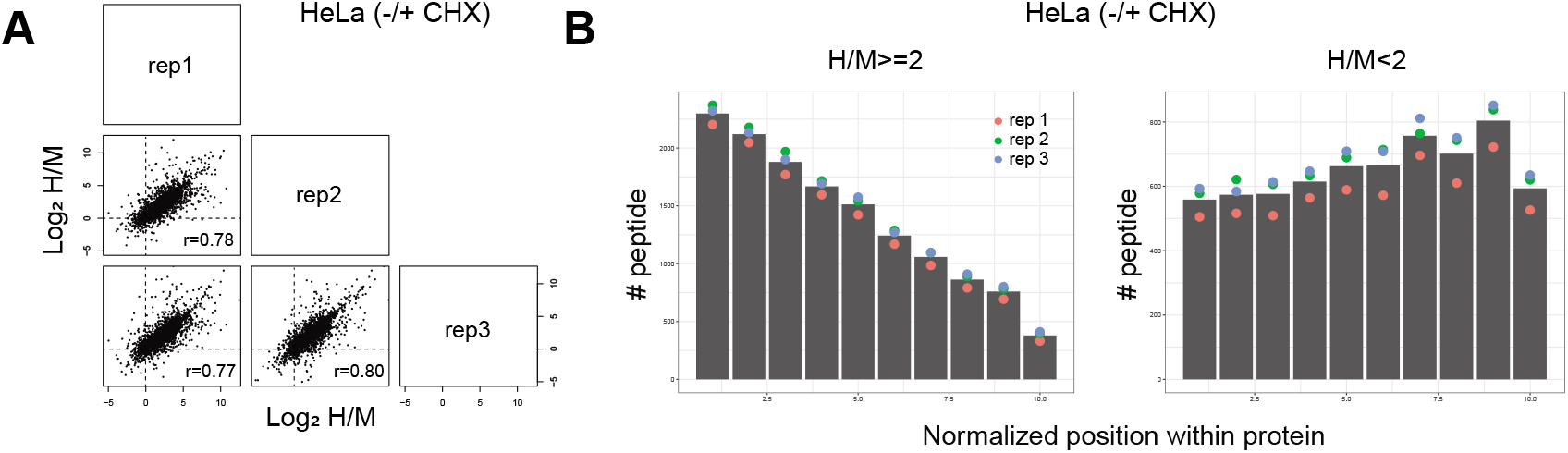
Quantitative analysis of nascent protein N-terminal acetylation. **(A)** Multi-scatter plots of log_2_ H/M ratios from three independent experiments using HeLa cells treated with DMSO or CHX. **(B)** pSNAP can enrich NPCs. Relative starting positions of identified peptides within proteins for enriched (left) or input (right) samples. The bars represent averaged values from three independent experiments.

## Acknowledgements

We are very grateful to Erik McShane (Harvard Medical School), Matthew L Kraushar (Charité-Universitätsmedizin Berlin), and Tatsuya Niwa (Tokyo Institute of Technology) for critical reading of the manuscript. We thank Takuya Uehata and Osamu Takeuchi for providing cultured cells. We thank the members of the Department of Molecular & Cellular BioAnalysis, the Department of Proteomics, Drug Discovery, Makoto Arita, Shinya Oki, Yosuke Isobe, Yasuhiro Murakawa, and Hiroshi Tsugawa for fruitful discussion. KI thanks the Samuro Kakiuchi Memorial Research Award for Young Scientists for supporting this study. This work was supported by JSPS Grant-in-Aid for Scientific Research (Grant Numbers 18K14674, 20H03241, 20H04844, 21H05720 to KI and 17H05667 to YI), JST PRESTO (JPMJPR18H2), JST FOREST (JPMJFR214L), JST ERATO (JPMJER2101), the Takeda Science Foundation to KI, AMED (18dm0307023h) to DOW, JST Strategic Basic Research Program CREST (18070870), and AMED Advanced Research and Development Programs for Medical Innovation CREST (18068699) to YI.

## Author contributions

Conceptualization & Methodology, J.U., Y.I., and K.I.; Investigation, J.U., R.R., K.M., Y.K., M.I., Y.M., D.O.W., Y.C, Y. I., and K.I; Resources, D.O.W., M.I, Y.M., Y.I., and K.I.; Writing - Original Draft, K.I; Writing - Review & Editing, J.U., R.R., K.M., M.I. Y.M, D.O.W., Y.I., Supervision, Y.I. and K.I..

## Competing interests

The authors declare no competing financial interest.

## Supplementary information

**Figures S1–4**

**Table S1 (related to Figure 1D)**: LC/MS/MS proteomic analysis of NPCs from HeLa cells treated with (H) or without (M) puromycin.

**Table S2 (related to Figures S2B and S2C)**: LC/MS/MS proteomic analysis of NPCs by pSNAP with short pulse labeling. LC/MS/MS proteomic analysis of NPCs from mouse RAW264.7 macrophages treated with LPS (H) or water (M).

**Table S3 (related to Figures 2A and 2D)**: LC/MS/MS proteomic analysis of NPCs from mouse primary cells.

**Table S4 (related to Figures 3B and 3D)**: LC/MS/MS differential proteomic analysis of NPCs from HeLa cells treated with DMSO (M) or silmitasertib (H). Statistical comparisons were made using two-sided Wilcoxon rank-sum test.

**Table S5 (related to Figures S4A and 4B)**: LC/MS/MS differential proteomic analysis of NPCs from HeLa cells treated with CHX (M) or DMSO (H). Unique protein Nt-acetylated sites that exhibit H/M≧2 are also shown.

**Table S6 (related to Figure 4G)**: LC/MS/MS differential proteomic analysis of peptide-based pulldown experiments on HeLa cells. Statistical comparisons were made using two-sided Student’s t-test.

## Highlights

- Nascent polypeptidome analysis with a simplified protocol.
- Quantification of acute changes in nascent polypeptides induced by external stimuli.
- Profiling and characterization of chemical modifications on nascent polypeptides.

## Notes

### Competing Interest Statement

The authors have declared no competing interest.

### Summary of Updates

Most of the main and supplementary figures revised.

## References

Aksnes, H., Ree, R., and Arnesen, T. (2019). Co-translational, Post-translational, and Non-catalytic Roles of N-Terminal Acetyltransferases. Mol. Cell 73, 1097–1114.

Aviner, R. (2020). The science of puromycin: From studies of ribosome function to applications in biotechnology. Comput. Struct. Biotechnol. J. 18, 1074–1083.

Aviner, R., Geiger, T., and Elroy-Stein, O. (2013). Novel proteomic approach (PUNCH-P) reveals cell cycle-specific fluctuations in mRNA translation. Genes Dev. 27, 1834–1844.

Aviner, R., Li, K.H., Frydman, J., and Andino, R. (2021). Cotranslational prolyl hydroxylation is essential for flavivirus biogenesis. Nature 596, 558–564.

Bertolini, M., Fenzl, K., Kats, I., Wruck, F., Tippmann, F., Schmitt, J., Auburger, J.J., Tans, S., Bukau, B., and Kramer, G. (2021). Interactions between nascent proteins translated by adjacent ribosomes drive homomer assembly. Science 371, 57–64.

Bouhaddou, M., Memon, D., Meyer, B., White, K.M., Rezelj, V.V., Correa Marrero, M., Polacco, B.J., Melnyk, J.E., Ulferts, S., Kaake, R.M., et al. (2020). The Global Phosphorylation Landscape of SARS-CoV-2 Infection. Cell 182, 685–712.e19.

Choudhary, C., Weinert, B.T., Nishida, Y., Verdin, E., and Mann, M. (2014). The growing landscape of lysine acetylation links metabolism and cell signalling. Nat. Rev. Mol. Cell Biol. 15, 536–550.

Collart, M.A., and Weiss, B. (2020). Ribosome pausing, a dangerous necessity for cotranslational events. Nucleic Acids Res. 48, 1043–1055.

Dieterich, D.C., Link, A.J., Graumann, J., Tirrell, D.A., and Schuman, E.M. (2006). Selective identification of newly synthesized proteins in mammalian cells using bioorthogonal noncanonical amino acid tagging (BONCAT). Proc. Natl. Acad. Sci. U. S. A. 103, 9482–9487.

Doherty, M.K., Hammond, D.E., Clague, M.J., Gaskell, S.J., and Beynon, R.J. (2009). Turnover of the human proteome: determination of protein intracellular stability by dynamic SILAC. J. Proteome Res. 8, 104–112.

Drazic, A., Aksnes, H., Marie, M., Boczkowska, M., Varland, S., Timmerman, E., Foyn, H., Glomnes, N., Rebowski, G., Impens, F., et al. (2018). NAA80 is actin’s N-terminal acetyltransferase and regulates cytoskeleton assembly and cell motility. Proc. Natl. Acad. Sci. U. S. A. 115, 4399–4404.

Eichelbaum, K., Winter, M., Berriel Diaz, M., Herzig, S., and Krijgsveld, J. (2012). Selective enrichment of newly synthesized proteins for quantitative secretome analysis. Nat. Biotechnol. 30, 984–990.

Forester, C.M., Zhao, Q., Phillips, N.J., Urisman, A., Chalkley, R.J., Oses-Prieto, J.A., Zhang, L., Ruggero, D., and Burlingame, A.L. (2018). Revealing nascent proteomics in signaling pathways and cell differentiation. Proc. Natl. Acad. Sci. U. S. A. 115, 2353–2358.

Frese, C.K., Mikhaylova, M., Stucchi, R., Gautier, V., Liu, Q., Mohammed, S., Heck, A.J.R., Altelaar, A.F.M., and Hoogenraad, C.C. (2017). Quantitative Map of Proteome Dynamics during Neuronal Differentiation. Cell Rep. 18, 1527–1542.

Gandin, V., Masvidal, L., Cargnello, M., Gyenis, L., McLaughlan, S., Cai, Y., Tenkerian, C., Morita, M., Balanathan, P., Jean-Jean, O., et al. (2016). mTORC1 and CK2 coordinate ternary and eIF4F complex assembly. Nat. Commun. 7, 11127.

Hansen, B.K., Gupta, R., Baldus, L., Lyon, D., Narita, T., Lammers, M., Choudhary, C., and Weinert, B.T. (2019). Analysis of human acetylation stoichiometry defines mechanistic constraints on protein regulation. Nat. Commun. 10, 1055.

Helbig, A.O., Gauci, S., Raijmakers, R., van Breukelen, B., Slijper, M., Mohammed, S., and Heck, A.J.R. (2010). Profiling of N-acetylated protein termini provides in-depth insights into the N-terminal nature of the proteome. Mol. Cell. Proteomics 9, 928–939.

Howden, A.J.M., Geoghegan, V., Katsch, K., Efstathiou, G., Bhushan, B., Boutureira, O., Thomas, B., Trudgian, D.C., Kessler, B.M., Dieterich, D.C., et al. (2013). QuaNCAT: quantitating proteome dynamics in primary cells. Nat. Methods 10, 343–346.

Hsieh, A.C., Liu, Y., Edlind, M.P., Ingolia, N.T., Janes, M.R., Sher, A., Shi, E.Y., Stumpf, C.R., Christensen, C., Bonham, M.J., et al. (2012). The translational landscape of mTOR signalling steers cancer initiation and metastasis. Nature 485, 55–61.

Huang, Y., Zhang, Q., Yang, L., Lin, L., Xie, J., Yao, J., Zhou, X., Zhang, L., Shen, H., and Yang, P. (2021). Puromycin-Modified Silica Microsphere-Based Nascent Proteomics Method for Rapid and Deep Nascent Proteome Profile. Anal. Chem. 93, 6403–6413.

Ingolia, N.T., Ghaemmaghami, S., Newman, J.R.S., and Weissman, J.S. (2009). Genome-wide analysis in vivo of translation with nucleotide resolution using ribosome profiling. Science 324, 218–223.

Kamenova, I., Mukherjee, P., Conic, S., Mueller, F., El-Saafin, F., Bardot, P., Garnier, J.-M., Dembele, D., Capponi, S., Timmers, H.T.M., et al. (2019). Co-translational assembly of mammalian nuclear multisubunit complexes. Nat. Commun. 10, 1740.

Keshwani, M.M., Klammt, C., von Daake, S., Ma, Y., Kornev, A.P., Choe, S., Insel, P.A., and Taylor, S.S. (2012). Cotranslational cis-phosphorylation of the COOH-terminal tail is a key priming step in the maturation of cAMP-dependent protein kinase. Proc. Natl. Acad. Sci. U. S. A. 109, E1221–E1229.

Kii, I., Sumida, Y., Goto, T., Sonamoto, R., Okuno, Y., Yoshida, S., Kato-Sumida, T., Koike, Y., Abe, M., Nonaka, Y., et al. (2016). Selective inhibition of the kinase DYRK1A by targeting its folding process. Nat. Commun. 7, 11391.

Klann, K., Tascher, G., and Münch, C. (2020). Functional Translatome Proteomics Reveal Converging and Dose-Dependent Regulation by mTORC1 and eIF2α. Mol. Cell 77, 913–925.e4.

Lai, Z.W., Petrera, A., and Schilling, O. (2015). Protein amino-terminal modifications and proteomic approaches for N-terminal profiling. Curr. Opin. Chem. Biol. 24, 71–79.

Lamper, A.M., Fleming, R.H., Ladd, K.M., and Lee, A.S.Y. (2020). A phosphorylation-regulated eIF3d translation switch mediates cellular adaptation to metabolic stress. Science 370, 853–856.

Marciano, R., Leprivier, G., and Rotblat, B. (2018). Puromycin labeling does not allow protein synthesis to be measured in energy-starved cells. Cell Death Dis. 9, 1–3.

McShane, E., Sin, C., Zauber, H., Wells, J.N., Donnelly, N., Wang, X., Hou, J., Chen, W., Storchova, Z., Marsh, J.A., et al. (2016). Kinetic Analysis of Protein Stability Reveals Age-Dependent Degradation. Cell 167, 803–815.e21.

Mellacheruvu, D., Wright, Z., Couzens, A.L., Lambert, J.-P., St-Denis, N.A., Li, T., Miteva, Y.V., Hauri, S., Sardiu, M.E., Low, T.Y., et al. (2013). The CRAPome: a contaminant repository for affinity purification-mass spectrometry data. Nat. Methods 10, 730–736.

Miyamoto-Sato, E., Nemoto, N., Kobayashi, K., and Yanagawa, H. (2000). Specific bonding of puromycin to full-length protein at the C-terminus. Nucleic Acids Res. 28, 1176–1182.

Mollapour, M., and Neckers, L. (2012). Post-translational modifications of Hsp90 and their contributions to chaperone regulation. Biochim. Biophys. Acta 1823, 648–655.

Mueller, F., Friese, A., Pathe, C., da Silva, R.C., Rodriguez, K.B., Musacchio, A., and Bange, T. (2021). Overlap of NatA and IAP substrates implicates N-terminal acetylation in protein stabilization. Sci Adv 7.

Olsen, J.V., Blagoev, B., Gnad, F., Macek, B., Kumar, C., Mortensen, P., and Mann, M. (2006). Global, in vivo, and site-specific phosphorylation dynamics in signaling networks. Cell 127, 635–648.

Pankow, S., Bamberger, C., and Yates, J.R., 3rd (2019). A posttranslational modification code for CFTR maturation is altered in cystic fibrosis. Sci. Signal. 12.

Parsons, J.L., Dianova, I.I., Finch, D., Tait, P.S., Ström, C.E., Helleday, T., and Dianov, G.L. (2010). XRCC1 phosphorylation by CK2 is required for its stability and efficient DNA repair. DNA Repair 9, 835–841.

Pisareva, V.P., Skabkin, M.A., Hellen, C.U.T., Pestova, T.V., and Pisarev, A.V. (2011). Dissociation by Pelota, Hbs1 and ABCE1 of mammalian vacant 80S ribosomes and stalled elongation complexes. EMBO J. 30, 1804–1817.

Reineke, L.C., Tsai, W.-C., Jain, A., Kaelber, J.T., Jung, S.Y., and Lloyd, R.E. (2017). Casein Kinase 2 Is Linked to Stress Granule Dynamics through Phosphorylation of the Stress Granule Nucleating Protein G3BP1. Mol. Cell. Biol. 37.

Schwanhäusser, B., Gossen, M., Dittmar, G., and Selbach, M. (2009). Global analysis of cellular protein translation by pulsed SILAC. Proteomics 9, 205–209.

Schwarz, A., and Beck, M. (2019). The Benefits of Cotranslational Assembly: A Structural Perspective. Trends Cell Biol. 29, 791–803.

Siddiqui-Jain, A., Drygin, D., Streiner, N., Chua, P., Pierre, F., O’Brien, S.E., Bliesath, J., Omori, M., Huser, N., Ho, C., et al. (2010). CX-4945, an orally bioavailable selective inhibitor of protein kinase CK2, inhibits prosurvival and angiogenic signaling and exhibits antitumor efficacy. Cancer Res. 70, 10288–10298.

Stumpf, C.R., Moreno, M.V., Olshen, A.B., Taylor, B.S., and Ruggero, D. (2013). The translational landscape of the mammalian cell cycle. Mol. Cell 52, 574–582.

Sugiyama, N., Imamura, H., and Ishihama, Y. (2019). Large-scale Discovery of Substrates of the Human Kinome. Sci. Rep. 9, 10503.

Thoreen, C.C., Chantranupong, L., Keys, H.R., Wang, T., Gray, N.S., and Sabatini, D.M. (2012). A unifying model for mTORC1-mediated regulation of mRNA translation. Nature 485, 109–113.

Timms, R.T., Zhang, Z., Rhee, D.Y., Harper, J.W., Koren, I., and Elledge, S.J. (2019). A glycine-specific N-degron pathway mediates the quality control of protein -myristoylation. Science 365.

Tong, M., Suttapitugsakul, S., and Wu, R. (2020). Effective Method for Accurate and Sensitive Quantitation of Rapid Changes of Newly Synthesized Proteins. Anal. Chem. 92, 10048–10057.

Uchiyama, J., Ishihama, Y., and Imami, K. (2021). Quantitative nascent proteome profiling by dual-pulse labelling with O-propargyl-puromycin and stable isotope-labelled amino acids. J. Biochem. 169, 227–236.

Varland, S., Osberg, C., and Arnesen, T. (2015). N-terminal modifications of cellular proteins: The enzymes involved, their substrate specificities and biological effects. Proteomics 15, 2385–2401.

Yeom, J., Ju, S., Choi, Y., Paek, E., and Lee, C. (2017). Comprehensive analysis of human protein N-termini enables assessment of various protein forms. Sci. Rep. 7, 6599.

