## Supplementary material for "pSNAP: Proteome-wide analysis of elongating nascent polypeptide chains": Method

### KEY RESOURCES TABLE

| REAGENT or RESOURCE | SOURCE | IDENTIFIER |
|---|---|---|
| **Antibodies** | | |
| mouse anti-puromycin antibody (clone 12D10) (IP) | Merck | Cat# MABE343, RRID:AB_2566826 |
| mouse anti-puromycin antibody (clone 3RH11) (1:5,000) | Cosmo Bio | Cat# CAC-PEN-MA001, RRID:AB_2620162 |
| rabbit anti-ubiquitin antibody (1:2,000) | Cell Signaling Technology | Cat# 3933, RRID:AB_2180538 |
| **Chemicals, peptides, and recombinant proteins** | | |
| puromycin dihydrochloride | FUJIFILM Wako | Cat# 160-23151 |
| L-($^{13}C_6$,$^{15}N_4$)-arginine (Arg”10”) | Cambridge Isotope Laboratories | Cat# CNLM-539-H |
| L-($^{13}C_6$,$^{15}N_2$)-lysine (Lys”8”) | Cambridge Isotope Laboratories | Cat# CNLM-291-H |
| L-($^{15}N_4$)-arginine (Arg”4”) | Cambridge Isotope Laboratories | Cat# LM-396 |
| L-($D_4$)-lysine (Lys”4”) | Cambridge Isotope Laboratories | Cat# DLM-2640 |
| MG-132 | Chemscene | Cat# CS-0471 |
| bortezomib | FUJIFILM Wako | Cat# 021-18901 |
| silmitasertib | MedChemExpress | Cat# HY-50855 |
| cycloheximide (CHX) | FUJIFILM Wako | Cat# 037-20991 |
| fetal bovine serum (FBS) | Thermo Fisher Scientific | Cat#10270106 |
| DMEM(High Glucose) with L-Glutamine, Phenol Red and Sodium Pyruvate | FUJIFILM Wako | Cat# 043-30085 |
| arginine- and lysine-free Neurobasal-A medium | Research Institute for the Functional Peptides | N/A |
| DMEM for SILAC | Thermo Fisher Scientific | Cat# 88364 |

| | | |
|---|---|---|
| neuron dissociation solutions | FUJIFILM Wako | Cat#291-78001 |
| poly-L-lysine | Merck | Cat#P6282 |
| MEM | Thermo Fisher Scientific | Cat#11095080 |
| D-glucose | Merck | Cat#A24940 |
| sodium pyruvate | Nacalai Tesque | Cat#06977-34 |
| penicillin-streptomycin | Thermo Fisher Scientific | Cat#15140122 |
| Neurobasal-A medium | Thermo Fisher Scientific | Cat#10888022 |
| B-27 supplement | Thermo Fisher Scientific | Cat#17504044 |
| GlutaMAX | Thermo Fisher Scientific | Cat#35050061 |
| LPS Salmonella minnesota R595 - TLR4 ligand | InvioGen | Cat#15C10-MM |
| bis(sulfosuccinimidyl)suberate, disodium salt (BS3) | FUJIFILM Wako | Cat# B574 |
| Sodium deoxycholate (SDC) | FUJIFILM Wako | Cat#194-08311 |
| Sodium N-lauroylsarcosinate (SLS) | FUJIFILM Wako | Cat#192-10382 |
| Trifluoroacetic acid (TFA) | FUJIFILM Wako | Cat#204-02743 |
| Ammonium acetate | FUJIFILM Wako | Cat#015-02837 |
| Dimethyl sulfoxide (DMSO) | FUJIFILM Wako | Cat#041-29351 |
| Ponceau-S Staining Solution | Beacle | Cat#BCL-PSS-01 |
| 100 x protease inhibitor cocktail | Merck | Cat#P8340 |
| Lysyl endopeptidase (LysC) | FUJIFILM Wako | Cat#129-02541 |
| Trypsin | Promega | Cat#V5111 |
| SETAPAETATPAPVEKSPAKK-K(biotin), Ac-SETAPAETATPAPVEKSPAKK-K(biotin), Ac-SETAPAETATPAPVEKpSPAKK-K(biotin) | Synpeptide | N/A |
| Nuclease-Free Water (not DEPC-Treated) | Thermo Fisher Scientific | Cat# AM9937 |
| TurboDNase (2U/uL) | Thermo Fisher Scientific | Cat# AM2238 |

| | | |
|---|---|---|
| SUPERaseIn | Thermo Fisher Scientific | Cat# AM2694 |
| cOmplete Mini, EDTA-free Protease Inhibitor Cocktail | Sigma-Aldrich | Cat# 11836170001 |
| QIAzol lysis reagent | QIAGEN | Cat# 79306 |
| Trizol LS reagent | Thermo Fisher Scientific | Cat# 10296028 |
| **Critical commercial assays** | | |
| SuperScript III First-Strand Synthesis SuperMix for qRT-PCR | Thermo Fisher Scientific | Cat# 11752050 |
| TaqMan Gene Expression Master Mix | Thermo Fisher Scientific | Cat# 4369016 |
| Empore SDB-XC membrane | GL Sciences | Cat#5010-30016 |
| Empore cation-SR membrane | GL Sciences | Cat#5010-30031 |
| Dynabeads Protein G Magnetic Beads | Thermo Fisher Scientific | Cat#10003D |
| Pierce™ High Capacity Streptavidin Agarose | Thermo Fisher Scientific | Cat#20357 |
| ReproSil-Pur C18-AQ materials 120 (3 μm) | Dr. Maisch GmbH, Ammerbuch | Cat#r13.aq. |
| ECL reagent | Cytiva | Cat#RPN2124 |
| **Deposited data** | | |
| Poteomic raw data | this paper | jPOST: PXD024078 |
| | | jPOST: PXD031475 |
| | | jPOST: PXD024080 |
| | | jPOST: PXD024081 |
| | | jPOST: PXD024104 |
| | | jPOST: PXD026349 |
| **Experimental models: Cell lines** | | |
| Human: HeLa | ATCC | Cat# CCL-2, RRID: CVCL_0030 |

| | | |
|---|---|---|
| Mouse: RAW264.7 | ATCC | Cat#TIB-71, RRID:CVCL_0493 |
| **Oligonucleotides** | | |
| RPS27 (Hs01378332_g1) TaqMan Assay | Thermo Fisher Scientific | Cat# 4331182 |
| RPL12 (Hs02385039_g1) TaqMan Assay | Thermo Fisher Scientific | Cat# 4331182 |
| RPL5 (Hs00851991_u1) TaqMan Assay | Thermo Fisher Scientific | Cat# 4331182 |
| RPS17 (Hs00734303_g1) TaqMan Assay | Thermo Fisher Scientific | Cat# 4331182 |
| GammaTub23C (Dm01841764_g1) TaqMan Assay | Thermo Fisher Scientific | Cat# 4331182 |
| **Software and algorithms** | | |
| MaxQuant (v1.6.17.0 or v.1.6.2.10) | Cox and Mann, 2008 | https://www.maxquant.org/ |
| Perseus (v.1.6.5.0) | Tyanova, S. *et al.* 2016 | https://maxquant.net/perseus/ |
| Metascape | Zhou, Y. *et al.* 2019 | https://metascape.org/gp/index.html#/main/step1 |
| FAIMS MzXML Generator | Hebert et al., 2018 | https://github.com/coongroup/FAIMS-MzXML-Generator |
| R Studio (v 1.2.5001) | | https://www.rstudio.com/ |

### EXPERIMENTAL MODEL AND SUBJECT DETAILS

#### HeLa cell culture and pulse labeling with puromycin and SILAC amino acids

HeLa cells (ATCC) were cultured in Dulbecco's modified Eagle's medium (DMEM) (FUJIFILM Wako) supplemented with 10% fetal bovine serum (FBS) (Thermo Fisher Scientific). Cells were grown to approximately 70–80% confluence and then used for experiments. For pulse-labeling experiments, the cell culture medium was switched to arginine- and lysine-free DMEM (Thermo Fisher Scientific) supplemented with 10% FBS and 10 μM puromycin (FUJIFILM Wako), either “heavy” amino acids [0.398 mM L-($^{13}C_6$,$^{15}N_4$)-arginine (Arg”10”) and 0.798 mM L-($^{13}C_6$,$^{15}N_2$)-lysine (Lys”8”)] or “medium” amino acids [0.398 mM L-($^{15}N_4$)-arginine (Arg”4”) and 0.798 mM L-($D_4$)-lysine (Lys”4”)] (Cambridge Isotope Laboratories), and incubated for 30 min or 2 hr as described elsewhere (Imami et al., 2010, 2018; Uchiyama et al., 2020). For the proof-of-concept experiment (related to **Figure 1B**), HeLa cells were pulse-labeled with a combination of 10 µM puromycin and ‘heavy’ amino acids (Arg’10’ and Lys’8’) for 2 hr, while as control, cells were treated with only ‘medium-heavy’ amino acids (Arg’4’ and Lys’4’) and puromycin was omitted. To examine the degradation of puromycylated NPCs during puromycin labeling (related to **Figure S1B**), HeLa cells were treated with 10 µM puromycin and either DMSO, 10 µM MG-132 (Chemscene) or 1 µM bortezomib (FUJIFILM Wako) for 15 min, 30 min, 60 min, and 120 min. In the silmitasertib experiment (related to **Figure 3A**), HeLa cells were pre-incubated with either DMSO or a CK2 inhibitor silmitasertib (MedChemExpress) (10 µM) for 10 min, and then pulse-labeled with puromycin for 2 hr in the presence of either DMSO (+‘medium-heavy’ amino acids) or 10 µM silmitasertib (+‘heavy’ amino acids). For the protein Nt-acetylation profiling (related to **Figure 4A**), HeLa cells were pulse-labeled with puromycin for 2 hr in the presence of either 100 µg/mL CHX (FUJIFILM Wako) (+‘medium-heavy’ amino acids) or DMSO (+‘heavy’ amino acids). After pulse-labeling, cells were washed twice with ice-cold PBS and collected by centrifugation. Three independent experiments were performed in all proteomic studies. All cells were maintained in an incubator at 37°C under humidified 5% $CO_2$ in air.

**Mouse primary cortex cultures and pulse labeling**

Primary cultures of cortical neurons were prepared as described with a few modifications (Kaech and Banker, 2006). In brief, cortices were dissected from postnatal day 0 (P0) mice. Cortical neurons were dissociated using neuron dissociation solutions (FUJIFILM Wako) and plated on 35 mm (DIV7 neuron pulse labeling) or 60 mm (DIV5 and DIV14 neuron pulse labelling) dishes coated with poly-L-lysine (Merck) at a density of $1x10^5$ and $2x10^5$ cells/$cm^2$, respectively, in MEM (Thermo Fisher Scientific) supplemented with 10% horse serum (Thermo Fisher Scientific), 0.6% D-glucose (Merck), 1 mM sodium pyruvate (Nacalai Tesque) and 1% penicillin-streptomycin (Thermo Fisher Scientific). Three hours after plating, the media was replaced by growth media consisting of Neurobasal-A medium (Thermo Fisher Scientific) supplemented with B-27 supplement (Thermo Fisher Scientific), GlutaMAX (Thermo Fisher Scientific) and penicillin-streptomycin (Thermo Fisher Scientific). Neurons were maintained at 37°C in 5% $CO_2$ until experiments. For pulse labeling experiments at 7 days in vitro (related to

**Figure 2A**) and experiments to assess differential translations between 5 and 14 days in vitro (related to **Figure 2D**), the cell culture medium was switched to arginine- and lysine-free Neurobasal-A medium (Research Institute for the Functional Peptides) supplemented with B-27 supplement, GlutaMAX, penicillin-streptomycin and either “heavy” amino acids or “medium-heavy” amino acids, and processed as described above. ICR mice were purchased (Shimizu). This study was carried out in accordance with the Guide for the Care and Use of Laboratory Animals from the Society for Neuroscience and was authorized by the Animal Care and Use Committee of Kyoto University.

**RAW264.7 cell culture and pulse labeling**

RAW264.7 cells (ATCC) were cultured in DMEM supplemented with 10% FBS. For lipopolysaccharide (LPS) stimulation experiments (related to **Figure S2C**), cells were grown to approximately 70% confluence and then used for experiments. Cells were pre-incubated with either 100 ng/mL LPS [Salmonella minnesota R595 - TLR4 ligand (InvivoGen 15C10-MM)] or vehicle (ultrapure water) for 1 hr, and then pulse-labeled with puromycin for 30 min, 1 hr or 2 hr in the presence of either ‘medium-heavy’ amino acids (for control cells) or ‘heavy’ amino acids (for LPS-stimulated cells). Three independent experiments were performed for each condition. Cells were maintained in an incubator at 37°C under humidified 5% $CO_2$ in air.

### METHOD DETAILS

**Immunoprecipitation (IP) of NPCs with anti-puromycin antibody**

HeLa cell pellets were lysed with a buffer [100 mM HEPES-NaOH (pH 7.5), 150 mM NaCl, 1% Nonidet P-40 (NP-40), protease inhibitor cocktail (Merck)], and cell debris was removed by centrifugation (4°C, 16,000 x g, 30 min). Mouse primary cells were washed twice with ice-cold PBS and directly lysed by using lysis buffer [100 mM HEPES-NaOH (pH 7.5), 150 mM NaCl, 1% Nonidet P-40 (NP-40), 1% protease inhibitor cocktail (Merck)] and further centrifuged at 21,500 x g for 5 min at 4°C to remove cell debris. The protein concentration was measured using a BCA assay (Thermo Fisher Scientific), and 250 µg (1 µg/µL) protein per sample was used for the following IP, except that 250 µg, 500 µg, or 750 µg protein was used for the experiment with 30 min pulse labeling (related to **Figure S2A**) and that 140 µg, 280 µg, and 560 µg protein were used for 2 hr, 1 hr, and 30 min pulse labeling in the LPS treatment experiments (related to **Figure S2C**). For the silmitasertib treatment experiments (related to **Figure 3A**), 125 µg protein inputs from ‘medium-heavy’- and ‘heavy’-labeled cells were combined (total 250 µg proteins). 62.6 µg Dynabeads™ Protein G Magnetic Beads (Thermo Fisher Scientific) and 15 µg anti-puromycin antibody (clone 12D10 from Merck Millipore or 3RH11 from Cosmo Bio) per IP experiment were mixed in PBS-0.02% Tween 20 (PBS-T) and incubated for 30 min at room temperature with rotation. The supernatant was removed and the beads were washed twice with a conjugation buffer (PBS). To crosslink the beads and the antibodies, the beads were suspended in 5 mM bis(sulfosuccinimidyl)suberate, disodium salt ($BS^3$, Wako FUJIFILM) in the conjugation buffer and incubated for 30 min at 37℃. To quench the reaction, 50 mM (final concentration) Tris-HCl pH 7.5 was added and incubation was continued for 15 min at room temperature. The antibody-conjugated beads were then rinsed with 0.02% PBS-T, PBS, and the lysis buffer. The antibody-conjugated beads were incubated with 250 µg protein input for 1 hr at 4°C with slow rotation. The supernatant was transferred to a new tube, and the beads were washed three times with PBS supplemented with 850 mM NaCl. Puromycylated NPCs were eluted from the beads with 100 µL 0.15% trifluoroacetic acid (TFA) (FUJIFILM Wako), and the elution was repeated once more. All TFA eluates were combined and dried in a SpeedVac (Thermo Fisher Scientific).

**Protein digestion and SCX fractionation**

The dried samples were resuspended with 100 mM Tris-HCl pH 9.0 containing 8 M urea. Proteins were reduced with 10 mM dithiothreitol (DTT) (FUJIFILM Wako) for 30 min at 37℃, followed by alkylation with 50 mM 2-iodoacetamide (IAA) (FUJIFILM Wako) for 30 min at room temperature in the dark. The samples were diluted to 2 M urea with 50 mM ammonium bicarbonate. The proteins were digested with 1 µg lysyl endopeptidase (LysC) (FUJIFILM Wako) and 1 µg trypsin (Promega) overnight at 37°C on a shaking incubator. The resulting peptides were acidified with 0.5% TFA (final concentration), and fractionated with a StageTip containing SDB-XC (upper) and SCX (bottom) Empore disk membranes (GL Sciences) (Adachi et al., 2016). Peptides were eluted from the tip sequentially using 1) 0.5% TFA and 30% acetonitrile (ACN) 2) 1% TFA and 30% ACN 3) 2% TFA and 30% ACN, 4) 3% TFA and 30% ACN 5) 3% TFA, 100 mM ammonium acetate and 30% ACN 6) 4% TFA, 500 mM ammonium acetate and 30% ACN, and 7) 500 mM ammonium acetate and 30% ACN (related to **Figures 1B, 2A, and 3A**). For protein Nt-acetylated profiling (related to **Figure 4A**), flowthrough and wash (with 0.1% TFA and 80% ACN) fractions, in which Nt-acetylated peptides are expected to be enriched were combined and measured by LC/MS/MS. The sample solution was evaporated in a SpeedVac and the residue was resuspended in 0.5% TFA and 4% ACN. For the experiments related to **Figures 2D, S2A, S2C, and 4G**, the digested peptides were desalted using SDB-XC StageTips prior to LC/MS/MS analysis (no SCX fractionation was performed).

**Peptide pulldown assay (related to Figure 4F)**

Synthetic peptides were purchased from Synpeptide, and peptide sequences used were as follows: SETAPAETATPAPVEKSPAKK-K(biotin), Ac-SETAPAETATPAPVEKSPAKK-K(biotin), and Ac-SETAPAETATPAPVEKpSPAKK-K(biotin). Synthetic peptides (60 nmol each) were incubated with 18 µL streptavidin agarose resins (high-capacity streptavidin agarose, Thermo Fisher Scientific) per experiment for 2 hr at room temperature in 80 µL of lysis buffer (1% NP-40, 150 mM NaCl, 25 mM Tris-HCl pH 7.5, and protease and phosphatase inhibitor cocktails). Synthetic peptides bound to agarose resins were incubated with HeLa cell lysate (300 µg in 500 uL lysis buffer) for 2 hr at 4°C. Resins were washed three times with 1 mL of lysis buffer, and proteins bound to resins were eluted with 60 µL of elution buffer [12 mM sodium deoxycholate, 12 mM sodium N-lauroylsarcosinate, 5 mM DTT, 100 mM Tris-HCl pH 9.0]. Afterward, samples were transferred to new tubes and processed for LC/MS/MS analysis as described elsewhere (Masuda et al., 2008)

**Mass spectrometry and data acquisition**

Nano-scale reversed-phase liquid chromatography coupled with tandem mass spectrometry (nanoLC/MS/MS) was performed by an Orbitrap Fusion Lumos mass spectrometer (Thermo Fisher Scientific) connected to a Thermo Ultimate 3000 RSLCnano pump and an HTC-PAL autosampler (CTC Analytics, Zwingen, Switzerland) equipped with a self-pulled analytical column (150 mm length × 100 µm i.d.) (Ishihama et al., 2002) packed with ReproSil-Pur C18-AQ materials (3 µm, Dr. Maisch GmbH, Ammerbuch, Germany). The mobile phases consisted of (A) 0.5% acetic acid and (B) 0.5% acetic acid and 80% ACN. Peptides were eluted from the analytical column at a flow rate of 500 nL/min with the following gradient: 5-10% B in 5 min, 10-40% B in 60 min, 40-99% B in 5 min, and 99% for 5 min.

For most of the experiments (related to **Figures 1B, 2A, 3A, and 4A**), the Orbitrap Fusion Lumos instrument was operated in the data-dependent mode with a full scan in the Orbitrap followed by MS/MS scans for 3 sec using higher-energy collisional dissociation (HCD). The applied voltage for ionization was 2.4 kV. The full scans were performed with a resolution of 120,000, a target value of $4x10^5$ ions, and a maximum injection time of 50 ms. The MS scan range was *m/z* 300–1,500. The MS/MS scans were performed with a 15,000 resolution, a $5x10^4$ target value, and a 50 ms maximum injection time. The isolation window was set to 1.6, and the normalized HCD collision energy was 30. Dynamic exclusion was applied for 20 sec. For the nascent proteome analyses of mouse primary cultures (**Figure 2D**) and RAW264.7 cells treated with LPS (**Figure S2C**), and the peptide pulldown assay (**Figure 4F**), the mass spectrometric analyses were carried out with the FAIMS Pro interface (Thermo Fisher Scientific). The compensation voltage (CV) was set to −40, −60, and -80, and the cycle time of each CV experiment was set to 1 s. The full scans were performed with a resolution of 120,000, the standard AGC target mode, and a maximum injection time of 50 ms. The MS scan range was *m/z* 300–1,500. The MS/MS scans were collected in the ion trap with the rapid mode, the standard AGC target mode, and a maximum injection time of 30 ms. The isolation window was set to 1.6, and the normalized HCD collision energy was 30. Dynamic exclusion was applied for 20 sec.

For the nascent proteome analyses with short pulse labeling (related to **Figure S2A**), the LC/MS/MS analyses were performed on an UltiMate 3000 RSLCnano system (Thermo Fisher Scientific), combined with an Orbitrap Exploris 480 mass spectrometer (Thermo Fisher Scientific). All mass spectrometric analyses were carried out in the data-dependent acquisition (DDA) mode. For the Orbitrap Exploris 480 system, peptides were separated on self-pulled needle columns (250 mm, 100 μm ID) packed with Reprosil-Pur 120 C18-AQ1.9 μm (Dr. Maisch, Ammerbuch, Germany) at 50 °C in a column oven (Sonation). The flow rate was 400 nL/min. The flow gradient was set as follows: 5% B in 5 min, 5−19% B in 55.3 min, 19−29% B in 21 min, 29−40% B in 8.7 min, and 40−99% B in 0.1 min, followed by 99% B for 4.9 min. The electrospray voltage was set to 2.4 kV in the positive mode. The mass spectrometric analysis was carried out with the FAIMS Pro interface. The FAIMS mode was set to a standard resolution, and the total carrier gas flow was 4.0 L/min. The CV was set to −40 and −60, and the cycle time of each CV experiment was set to 1 s. The mass range of the survey scan was from 375 to 1500 m/z with a resolution of 60,000, 300% normalized automatic gain control (AGC) target and auto maximum injection time. The first mass of the MS/MS scan was set to 120 m/z with a resolution of 15,000, standard AGC, and auto maximum injection time. Fragmentation was performed by HCD with a normalized collision energy of 30%. The dynamic exclusion time was set to 20 s.

**Processing of mass spectrometry data**

All raw data files were analyzed and processed by MaxQuant (Cox and Mann, 2008) (v1.6.17.0 or v.1.6.2.10), and the database search was performed with Andromeda(Cox et al., 2011) against the SwissProt database (version 2020-10, 42,372 human protein entries) or mouse UniProt database (version 2020-3, 55,462 protein entries) spiked with common contaminants and enzyme sequences. Raw data files collected from FAIMS experiments were split into a set of MaxQuant compliant MzXML files using FAIMS MzXML Generator (https://github.com/coongroup/FAIMS-MzXML-Generator) (Hebert et al., 2018). Search parameters included two missed cleavage sites and variable modifications such as L-($^{13}C_6$,$^{15}N_4$)-arginine (Arg10 = heavy Arg, +10.00827), L-($^{13}C_6$,$^{15}N_2$)-lysine (Lys8 = heavy Lys, +8.01420), L-($^{15}N_4$)-arginine (Arg 4 = medium-heavy Arg, +3.98814 Da), L-($D_4$)-lysine (Lys 4 = medium-heavy Lys, +4.02511), methionine oxidation, protein N-terminal acetylation and phosphorylation of tyrosine, serine and threonine (only for the silmitasertib experiment related to **Figure 3A** and Nt-acetylome experiment related to **Figure 4A**). Cysteine carbamidomethylation was set as a fixed modification. The peptide mass tolerance was 4.5 ppm, and the MS/MS tolerance was 20 ppm. The false discovery rate (FDR) was set to 1% at the peptide spectrum match (PSM) level and protein level.

For protein-level quantification in all experiments, ‘unique + razor’ peptides were used. For the SILAC-based protein quantification, a minimum of one ratio count (unique peptide ion) was used for quantification, and the ‘re-quantify’ and ‘match between runs’ functions were employed. Raw H/M ratios were used for quantification related to **Figures 1B**, **2A, 4A, and S2A** while normalized H/M ratios were used for the differential analysis of NPC levels (related to **Figures 2D, 3A, and S2C**). For label-free quantification of peptide pulldown samples (related to **Figure 4F**), a minimum of two ratio counts was used for quantification. Only proteins quantified in 2 out of the 3 replicates in at least one condition were used for further analysis, and missing values were imputed from a normal distribution of $\log_2$ LFQ intensity using a default setting (width 0.3, down shift 1.8) in Perseus (v.1.6.5.0) (Tyanova et al., 2016). Volcano plots were generated based on $\log_2$ FC (x-axis) and $-\log_{10}$ p-value from two-sided t-test (y-axis). The curve indicates a cut-off for differentially interacting proteins (FDR<0.05, s0: 0.8).

For the differential NPC analysis in mouse primary cortex cultures (related to **Figure 2D**), normalized H/M ratios were $\log_2$ transformed and replicates were averaged when they were quantified in at least two of the three replicates. Two-sided one sample t-tests were performed on the experimental data using Perseus (Tyanova et al., 2016) and proteins were considered as 'significantly regulated' when they were below a t-test p-value of 0.05 and above 0.5 (a $\log_2$ ratio) according to an FDR estimation of SILAC fold-change (Bogdanow et al., 2020). To do this, we used the simulated data as a false positive set and accepted proteins above a SILAC fold-change that recalled <5% false positives. Our criteria ( p-value of 0.05 and above 0.5 (a $\log_2$ ratio)) correspond to FDR=0.04. GO enrichment (**Figure 2G**) of the significantly regulated proteins were performed using Metascape (Zhou et al., 2019). Adjusted p-values were shown.

For phosphosite analysis (related to **Figure 3D**), only class I (localization probability>0.75) sites (Olsen et al., 2006) that were quantified in at least one of the replicates were used.

#### Analysis of the positions of identified peptides within protein sequences

To calculate the positions of identified peptides within protein sequences (related to **Figures 1E, 2C, S2B, S3A, and S4B**), we classified peptides into two groups based on their H/M ratios; H/M≧2 (i.e., NPCs) and H/M<2 (i.e., potential contaminants). The first amino acids within peptides were mapped onto a protein sequence, and their relative positions within proteins (from N-term: 0 to C-term: 1) were calculated. For the input samples, all identified peptides were used to compute the positions within proteins. Only M- and H-labeled peptides were considered to calculate the positions within protein sequences in **Figure S3A**.

#### Comparison of pSNAP with pSILAC and ribo-seq (related to Figure S1G)

To compare the three orthogonal approaches (pSNAP, pSILAC, and ribo-seq), we used intensity-based absolute quantification (iBAQ) intensities (Schwanhäusser et al., 2011) from proteomic analysis and read counts of ribosome protected fragments (RPFs) obtained from ribo-seq. The iBAQ algorithm computes the sum of all peptides intensities divided by the number of theoretically observable peptides, which provides a rough estimation of protein abundance within a sample. In pSNAP and pSILAC experiments (**Figure 1B**), iBAQ intensities from the 'heavy' channel were used as a relative protein abundance of nascent proteins within the sample and compared to the read counts of RPFs obtained from the ribo-seq. The pSILAC data was obtained from the input sample (**Figure 1E**: without enrichment of NPCs) in which 2 hr pulse labeling was performed. Ribo-seq data were obtained from a previously published data set (Stumpf et al., 2013) in which cell cycle-dependent (i.e., G1, S, M) translational changes were analysed in HeLa cells. Transcripts having read counts within 0–10 were eliminated. For our pSNAP result, we only used proteins showing H/M ratio > 2 as those proteins are more likely to be *bona fide* NPCs. To compare our proteomic result (asynchronous HeLa cells) with the ribosome profiling data, we used data from the G1 cell-cycle stage because G1 is the major cell-cycle phase in asynchronous cells.

**Western blotting**

The cell lysates were re-suspended in LiDS loading sample buffer (Thermo Fisher Scientific) with 50 mM DTT and incubated at 70ºC for 5 min. The protein samples were loaded onto a 4–12% gradient SDS-polyacrylamide gel (Thermo Fisher Scientific) and separated using electrophoresis. The proteins were then transferred to a PVDF membrane (Merck Millipore) using a semi-dry western blot transfer system set to a constant current of 200 mA for 30 min. The membranes were first blocked by incubation in 5% (w/v) BSA or 5% (w/v) skim milk in Tris-buffered saline and 0.1% tween 20 (TBS-T) and then incubated with anti-puromycin antibody (clone 3RH11, Cosmo Bio) diluted 1:5,000, overnight at room temperature. The membrane was washed three times with 0.1% TBS-T, incubated with HRP-conjugated anti-mouse secondary antibody (1: 20,000 dilution) in 0.1% TBS-T 1 hr at room temperature, washed three times in TBS-T, and developed with ECL reagent (Cytiva). Ubiquitinated proteins were detected using anti-ubiquitin rabbit antibody (CST) (1: 2,000 dilution) and HRP-conjugated anti-rabbit secondary antibody (1: 10,000 dilution). Chemiluminescence was detected using the luminescent image analyzer ImageQuant LAS-500 (Cytiva).

**qRT-PCR analysis of selected mRNAs in monosome and polysome fractions (related to Figure 3C)**

HeLa cells (70% confluence) were treated with either 10 μM silmitasertib or DMSO for 2 hr. Two independent experiments were performed. The cells were placed on ice and gently washed once with 10 mL ice-cold PBS. Then they were lysed with 0.4 mL ice-cold lysis buffer (20 mM Tris-HCl, pH 7.5, 150 mM NaCl, 5 mM $MgCl_2$, 1 mM DTT, Complete EDTA-free Protease Inhibitor Cocktail, 100 μg/mL cycloheximide, 1% Triton X-100, 25 units/mL Turbo DNase

(Thermo Fisher Scientific), and 100 units/mL SUPERaseIn (Thermo Fisher Scientific)). The lysate was incubated on ice for 10 min and triturated through a 25-gauge needle (Terumo) ten times before centrifugation at 20,000 x g for 10 min at 4°C. The supernatant was collected in a new tube for the sucrose gradient analysis and as the sample for loading. A 10%–45% continuous sucrose gradient was prepared in a polyclear tube (Seton) using 10% and 45% sucrose buffers containing 100 μg/mL cycloheximide and 1 mM DTT in polysome buffer (25 mM Tris-HCl (pH 7.5), 150 mM NaCl and 15 mM $MgCl_2$) and the Biocomp Gradient Master program (Biocomp). An equal amount of cell lysate to the sample (300 μL) was loaded on the prepared gradient solution. Monosome and polysomes were separated in the sucrose gradient by ultracentrifugation using a SW-41 rotor (Beckman Coulter) at 36,000 rpm for 2.5 h at 4°C.The profile of relative RNA abundances of monosomes and polysomes were visualized at 254-nm wavelength, and equal-volume fractions were collected simultaneously with the Biocomp Piston Gradient Fractionator (Biocomp).

For the RNA analysis, an equal sample volume of TRIzol LS reagent (Thermo Fisher Scientific) was immediately added to the fractions and loading sample. RNA was purified using an miRNeasy Mini Kit according to manufacturer's instructions. Purified RNAs along with 1 ng of spiked *Drosophila* RNA were used for single strand complementary DNA (cDNA) synthesis using a SuperScript III First-Strand Synthesis SuperMix for qRT-PCR (Thermo Fisher Scientific). Quantitative RT-PCR was performed using TaqMan Gene Expression Master Mix (Thermo Fisher Scientific) on a Quant Studio 3 instrument (Thermo Fisher Scientific). The cDNA synthesis rate is validated by the spiked *Drosophila* RNA and the sample Ct values were compared with the loading sample before the sucrose gradient.

### Quantification and statistical analysis

The type of statistical test (e.g., Wilcoxon rank-sum) is annotated in the Figure legend and/or in the Methods specific to the analysis. In addition, statistical parameters such as the value of n, mean/median, SD and significance level are reported in the Figures and/or in the Figure Legends. Statistical analyses were performed using R as described in Methods and Resources for each individual analysis.

### Data availability

The proteomics data have been deposited to the ProteomeXchange Consortium via the jPOST (Moriya et al., 2019; Okuda et al., 2017) partner repository with the dataset identifier, PXD024078 (pSNAP evaluation related to **Figures 1D and S2A**, URL https://repository.jpostdb.org/preview/3537547806200eaf5d9028 Access key 6331), PXD031475 (LPS experiment related to **Figure S2C**, URL https://repository.jpostdb.org/preview/11358768266200e838672df Access key 4242), PXD024080 (silmitasertib experiment related to **Figure 3A**, URL https://repository.jpostdb.org/preview/1825726826602c7a2ddaa8e Access key 5013), PXD024081 (Nt-acetylation related to **Figure 4A**, URL https://repository.jpostdb.org/preview/469670382602c7a50ac18e Access key 5636), PXD024104 (mouse primary culture related to **Figures 2A and 2D**, URL https://repository.jpostdb.org/preview/1381455481602c7a62a3f2f Access key 1344), \PXD026349 (peptide pulldown related to **Figure 4G**, URL https://repository.jpostdb.org/preview/112165041260b49083b0dda Access key 8894).

### References


Adachi, J., Hashiguchi, K., Nagano, M., Sato, M., Sato, A., Fukamizu, K., Ishihama, Y., and Tomonaga, T. (2016). Improved Proteome and Phosphoproteome Analysis on a Cation Exchanger by a Combined Acid and Salt Gradient. Anal. Chem. *88*, 7899–7903.

Bogdanow, B., Schmidt, M., Weisbach, H., Gruska, I., Vetter, B., Imami, K., Ostermann, E., Brune, W., Selbach, M., Hagemeier, C., et al. (2020). Cross-regulation of viral kinases with cyclin A secures shutoff of host DNA synthesis. Nat. Commun. *11*, 4845.

Cox, J., and Mann, M. (2008). MaxQuant enables high peptide identification rates, individualized p.p.b.-range mass accuracies and proteome-wide protein quantification. Nat. Biotechnol. *26*, 1367–1372.

Cox, J., Neuhauser, N., Michalski, A., Scheltema, R.A., Olsen, J.V., and Mann, M. (2011). Andromeda: a peptide search engine integrated into the MaxQuant environment. J. Proteome Res. *10*, 1794–1805.

Hebert, A.S., Prasad, S., Belford, M.W., Bailey, D.J., McAlister, G.C., Abbatiello, S.E., Huguet, R., Wouters, E.R., Dunyach, J.-J., Brademan, D.R., et al. (2018). Comprehensive Single-Shot Proteomics with FAIMS on a Hybrid Orbitrap Mass Spectrometer. Anal. Chem. *90*, 9529–9537.

Imami, K., Sugiyama, N., Tomita, M., and Ishihama, Y. (2010). Quantitative proteome and phosphoproteome analyses of cultured cells based on SILAC labeling without requirement of serum dialysis. Mol. Biosyst. *6*, 594–602.

Imami, K., Milek, M., Bogdanow, B., Yasuda, T., Kastelic, N., Zauber, H., Ishihama, Y.,

Landthaler, M., and Selbach, M. (2018). Phosphorylation of the Ribosomal Protein RPL12/uL11 Affects Translation during Mitosis. Mol. Cell *72*, 84–98.e9.

Ishihama, Y., Rappsilber, J., Andersen, J.S., and Mann, M. (2002). Microcolumns with self-assembled particle frits for proteomics. J. Chromatogr. A *979*, 233–239.

Kaech, S., and Banker, G. (2006). Culturing hippocampal neurons. Nat. Protoc. *1*, 2406–2415.

Masuda, T., Tomita, M., and Ishihama, Y. (2008). Phase transfer surfactant-aided trypsin digestion for membrane proteome analysis. J. Proteome Res. *7*, 731–740.

Moriya, Y., Kawano, S., Okuda, S., Watanabe, Y., Matsumoto, M., Takami, T., Kobayashi, D., Yamanouchi, Y., Araki, N., Yoshizawa, A.C., et al. (2019). The jPOST environment: an integrated proteomics data repository and database. Nucleic Acids Res. *47*, D1218–D1224.

Okuda, S., Watanabe, Y., Moriya, Y., Kawano, S., Yamamoto, T., Matsumoto, M., Takami, T., Kobayashi, D., Araki, N., Yoshizawa, A.C., et al. (2017). jPOSTrepo: an international standard data repository for proteomes. Nucleic Acids Res. *45*, D1107–D1111.

Olsen, J.V., Blagoev, B., Gnad, F., Macek, B., Kumar, C., Mortensen, P., and Mann, M. (2006). Global, in vivo, and site-specific phosphorylation dynamics in signaling networks. Cell *127*, 635–648.

Schwanhäusser, B., Busse, D., Li, N., Dittmar, G., Schuchhardt, J., Wolf, J., Chen, W., and Selbach, M. (2011). Global quantification of mammalian gene expression control. Nature *473*, 337–342.

Stumpf, C.R., Moreno, M.V., Olshen, A.B., Taylor, B.S., and Ruggero, D. (2013). The translational landscape of the mammalian cell cycle. Mol. Cell *52*, 574–582.

Tyanova, S., Temu, T., Sinitcyn, P., Carlson, A., Hein, M.Y., Geiger, T., Mann, M., and Cox, J. (2016). The Perseus computational platform for comprehensive analysis of (prote)omics data. Nat. Methods *13*, 731–740.

Uchiyama, J., Ishihama, Y., and Imami, K. (2020). Quantitative nascent proteome profiling by dual pulse labeling with O-propargyl-puromycin and stable isotope labeled amino acids. J. Biochem.

Zhou, Y., Zhou, B., Pache, L., Chang, M., Khodabakhshi, A.H., Tanaseichuk, O., Benner, C., and Chanda, S.K. (2019). Metascape provides a biologist-oriented resource for the analysis of systems-level datasets. Nat. Commun. *10*, 1523.
